# *Toxoplasma gondii* RAD51 recombinase is required to overcome DNA replication stress and its inactivation leads to bradyzoite differentiation

**DOI:** 10.1101/2025.04.08.647840

**Authors:** Ana M. Saldarriaga Cartagena, Ayelén Aparicio Arias, Constanza Cristaldi, Agustina Ganuza, M. Micaela Gonzalez, María M. Corvi, William J. Sullivan, Laura Vanagas, Sergio O. Angel

## Abstract

*Toxoplasma gondii* is an obligate intracellular parasite with a high replication rate that can lead to DNA replicative stress, in turn associated with the generation of DNA double-strand breaks (DSBs). Cells have two main pathways to repair DSBs: non-homologous end joining and homologous recombination repair (NHEJ and HRR respectively). RAD51 is the key recombinase in the HRR pathway. In this work, we achieved endogenous tagging of the RAD51 gene using the Auxin Inducible Degron (AID) system, to generate the clonal line RH RAD51^HA-AID^. Here we demonstrate that RAD51 is expressed in replicative tachyzoites and establishes damage foci. Auxin-induced knock-down (KD) affects the correct replication of tachyzoites which show loss of synchronization. The use of the RAD51 inhibitor B02 also affects parasite growth, with an IC_50_ of 4.8 µM. B02 produced alterations in tachyzoite replication and arrest in the S phase of the cell cycle. Additionally, B02 induced tachyzoite to bradyzoite differentiation showing small cyst-like structures. In conclusion, the HRR pathway is necessary for maintaining proper tachyzoite replication under normal growth conditions, supporting that replicative stress occurs during the cell cycle. Our findings also suggest that DNA replication stress can induce bradyzoite differentiation.

## Introduction

*Toxoplasma gondii* is an intracellular parasite that undergoes both sexual and asexual phases during its life cycle. The sexual phase occurs exclusively in the intestines of felines, where infectious oocysts are subsequently excreted into the environment. In contrast, the asexual phase — consisting of rapidly replicating tachyzoites and dormant bradyzoites — takes place in intermediate hosts, including birds, mammals, and humans [1]. The infection is contracted orally, either through ingestion of oocysts contaminating soil or water sources, or through tissue cysts present in raw or undercooked meats [2–4]. This protozoan parasite is responsible for the widespread infection known as human toxoplasmosis, affecting one-third of the population and producing opportunistic disease. Congenital toxoplasmosis can lead to spontaneous abortion or birth defects. Chronic toxoplasmosis can be fatal in AIDS/HIV patients [5].

During acute infection, tachyzoites can infect any nucleated cell, spreading rapidly throughout the body until the immune system controls *T. gondii* dissemination. At this point, *T. gondii* tachyzoites convert into bradyzoites, generating tissue cysts that lead to chronic infection, recently associated with neurological disorders [6, 7]. Currently, sulfadiazine and pyrimethamine are typically used for the treatment of acute toxoplasmosis, but they are associated with dermatologic and bone marrow suppression as secondary effects. There is no treatment for the chronic stage of infection [8]. To identify newer and safer drugs for the effective treatment of toxoplasmosis, it is crucial to investigate key aspects of the tachyzoite infection process to uncover potential vulnerabilities.

*T. gondii* replicates within the parasitophorous vacuole (PV) through a process named endodyogeny, which is synchronized, doubling the number of tachyzoites in a sequence: from 1 to 2, then 4, 8, and so forth [9, 10]. Among the most notable aspects of toxoplasmic infection is the high speed at which tachyzoite replication occurs, between 5 to 9 hours, depending on the strain [11]. The high rate of rapid cellular division is likely to induce significant DNA replication stress that the parasite must manage to maintain genomic stability.

DNA replication stress is a process that can activate replication restart mechanisms associated with homologous recombination repair (HRR) to resume replication fork progression and preserve genome integrity [12]. DNA replication stress can be induced by a lack of nucleotide availability, mismatches or DNA lesions leading to replication fork stalling as well as the activation of checkpoints that arrest the cell cycle. The persistence of fork stalling can lead to double strand breaks that triggers the HRR response [13].

Several components of standard DNA repair pathways are conserved in *T. gondii*, including the DSB repair response, however they remain largely understudied [14]. DSB is repaired mainly by HRR and Non-Homologous End Joining (NHEJ), although other alternative pathways, like single strand annealing (SSA) or alternative end joining (ALT-EJ) exist [15, 16]. The restriction of HRR to the late S-phase is linked to the requirement of homologous sequences as templates for DNA repair. *T. gondii* presents many components of the HHR and NHEJ pathways, although those associated with modulating the choice of pathways have not been detected [17]. Interestingly, most of the components of the HRR are essential in the lytic cycle [17, 18].

RAD51 recombinase is a key protein of the HRR mechanism [19]. Also, it is implicated in the template switching pathway, a single strand DNA repair mechanism during DNA replication [20]. RAD51 has an ATP-dependent DNA-binding activity site that multimerizes on single-stranded DNA to form a helical filament. Achanta and collaborators [21] characterized RAD51 from *T. gondii* (TgRAD51) and demonstrated that compromised ATPase activity of TgRAD51 leads to inefficient gene targeting and poor gene conversion efficiency. Importantly, evidence supports that RAD51 is a druggable target. Compound B02 was identified as an inhibitor of human RAD51 in a screen of over 200,000 compounds [22]. Its mechanism of action is to disrupt the binding of RAD51 to DNA [23]. It was also reported that B02 inhibits the ATPase activity of RAD51 in *Plasmodium falciparum* [24], another apicomplexan parasite. B02 blocks PfRAD51 self-interaction, preventing its assembly at DNA damage sites, which impairs DNA repair mechanisms and increases the susceptibility to genotoxic agents.

In this work, an inducible RAD51 knockdown (KD) mutant line (RAD51^HA-AID^), in which *T. gondii* RAD51 (TgRAD51) protein is tagged with 3 hemagglutinin (HA) epitopes and Auxin Inducible Degron (AID), was generated. The presence of TgRAD51 foci was observed in tachyzoites grown in normal conditions without genotoxic agents. Knockdown of TgRAD51 was not lethal but altered tachyzoite replication. Moreover, treatment of tachyzoites with B02 altered growth and cell cycle progression. B02 also facilitated spontaneous bradyzoite cyst formation. Overall, the results demonstrate that TgRAD51 plays a crucial role in resolving DNA replication stress, enabling cell cycle progression in *T. gondii*.

## Results

### Identification of the *T. gondii* RAD51 gene

There are 4 genes in *T. gondii* that could encode proteins with a possible recombinase function (Fig. S2). TGME49_272900 was suggested as the orthologous RAD51 [17]. This gene has the length, domains and S-phase expression expected for a RAD51. TGME49_321430 has a small region compatible with a recombinase. Its expression is more associated with G1 (ToxoDB transcriptome). TGME49_313710 has a RAMA domain, associated with an archaeal RecA [25]. Its expression is detectable in tachyzoites but increases in bradyzoites at 48 hours (ToxoDB transcriptome; [26]). TGME49_216400 corresponds to a DMC1 recombinase, which specializes in recombination during meiosis [27]. In *T. gondii*, TgDMC1 is expressed at very low levels during the tachyzoite stage throughout the cell cycle. For all these reasons, the gene TGME49_272900 was chosen as the only ortholog for TgRAD51.

### Establishment of the RAD5^HA-AID^ cell line

To study the relevance of TgRAD51 in the lytic cycle of tachyzoite, we generated a line of *T. gondii* expressing endogenous TgRAD51 protein tagged with AID plus 3 HA epitopes at its C-terminus (*T. gondii* RAD51^HA-AID^) (Fig. S1). TgRAD51-degradation can be induced by IAA. Parental TIR1 tachyzoites, in which this line was generated, is RHΔKU80 [28, 29]. RHΔKU80 is unable to repair DSB by NHEJ pathway due to the lack of Ku80 protein, relying only on the HRR pathway [30]. Therefore, in presence of auxin, RAD51^HA-AID^ tachyzoites would have both NHEJ and HRR repair mechanisms disabled, allowing us to understand the importance of Rad51 for HRR pathway.

The subcellular localization of TgRAD51 was analyzed in RAD51^HA-AID^ tachyzoites using anti-HA antibodies. Under normal growth conditions, TgRAD51 was detected as distinct foci in the nucleus (Fig. 1A). In the presence of IAA, TgRAD51 protein is degraded and foci disappear (Fig. 1B, C). Foci were not observed in the Parental line (Fig. S3). Therefore, these results indicate that during tachyzoite replication under normal conditions DNA replication stress events would appear.

**Fig. 1.**
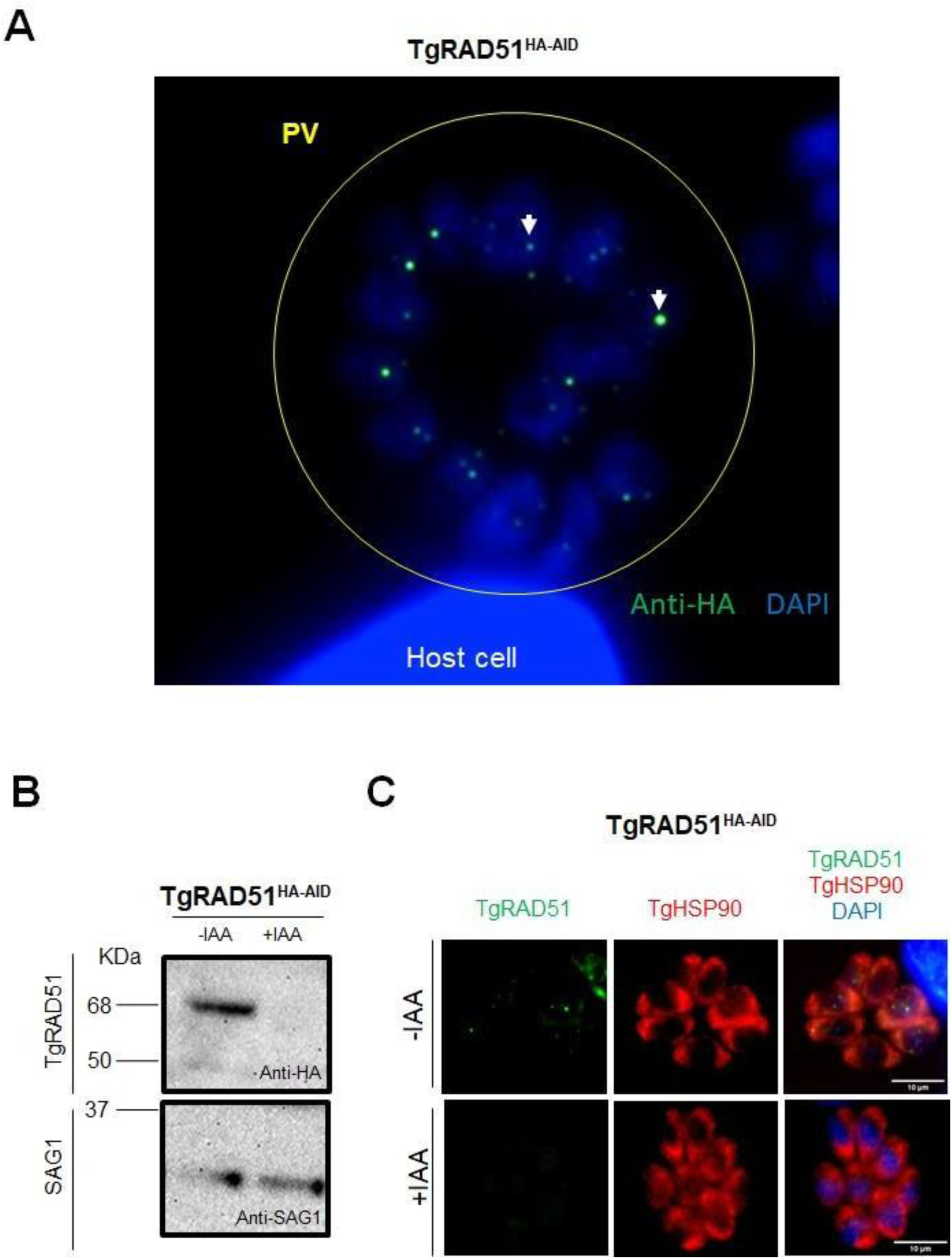
Generation of the *T. gondii* cell line RAD51^HA-AID^. The parental RHΔku80::Tir1 cell line was used to generate a *T. gondii* cell line expressing TgRAD51 fused to the auxin-inducible degron (AID) domain plus 3 hemagglutinin (HA) motifs. The protein expressed by the RAD51^HA-AID^ cell line was called TgRAD51. **A.** hTERT monolayers were infected with RAD51^HA-AID^ and analyzed by epifluorescence microscopy using anti-HA antibodies (green) and 4′,6-diamidino-2-phenylindole (DAPI). TgRAD51 foci are clearly visible in the *T. gondii* nucleus. PV: parasitophorous vacuole. White arrows show examples of TgRAD51 foci. **B.** Western blot of tachyzoites extracted from hTERT cultures infected with RAD51^HA-AID^ in the presence of DMSO or 500 μM Indole-3-acetic acid (IAA [auxin]) for 4 h. Anti-HA antibody was used to detect TgRAD51. Anti-SAG1 (Surface antigen 1) antibody was used as a loading control. The disappearance of TgRAD51 is observed in the presence of IAA after 4 h. **C.** Epifluorescence microscopy analysis of intracellular tachyzoites treated as in (**B**). DAPI and anti-HA and anti-*T. gondii* Hsp90 (TgHSP90, cytosolic) antibodies were used. The disappearance of TgRAD51 foci is observed after 4 h in the presence of IAA.

### TgRAD51 foci are associated with DNA replication stress

The presence of TgRAD51 foci was clearly observed in the absence of any external DNA damage agent. We reasoned that an internal source of DNA damage, such as DNA replication stress, could induce the expression of RAD51 and the generation of foci. To confirm this hypothesis, extracellular non-replicating RAD51^HA-AID^ tachyzoites were tested with anti-HA antibody. As shown in Figure 2 (A), no foci were detected in any tachyzoite, indicating that the ability to replicate is necessary for the appearance of RAD51.

**Fig. 2.**
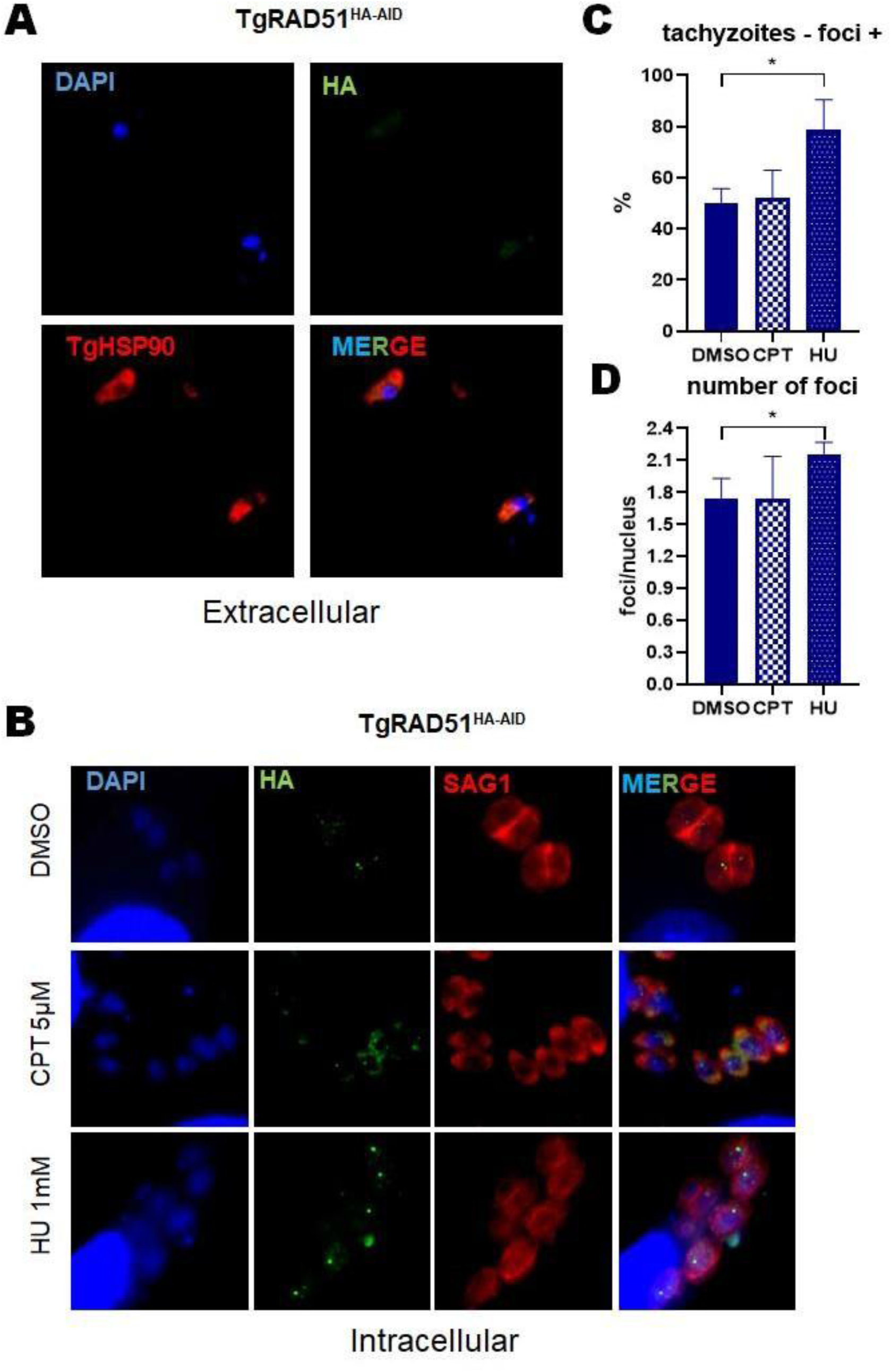
Analysis of DNA replication stress by TgRAD51 foci formation. **A.** Extracellular RAD51^HA-AID^ tachyzoites were labeled with anti-HA to evaluate foci formation. Anti-TgHSP90 antibody was used to label tachyzoite cytoplasm. **B**. hTERT monolayers infected with RAD51^HA-AID^ tachyzoites were treated with 1 mM HU, 4 µM CPT or vehicle (DMSO) for 4 hours and labeled with anti-HA to assess foci formation. Anti-SAG1 antibody was used to detect tachyzoites. **C**. The percentage of tachyzoites displaying foci in the presence of the genotoxic agents HU, CPT and DMSO was plotted. Data were analyzed by one-way ANOVA and Tukey’s multiple comparison test. *p<0.05. The graph is an example of three assays with similar results **D.** The number of TgRAD51 foci per tachyzoite was obtained in parasites treated with HU, CPT and DMSO. A total of 15 cores with foci selected from at least 5 randomly taken fields were analyzed. Data were analyzed by one-way ANOVA and Tukey’s multiple comparison test. *p<0.05. The graph is an example of three assays with similar results

If the presence of TgRAD51 foci were associated with DNA replication stress, they should increase in the presence of genotoxic agents that induce replication-associated DNA damage. Intracellular RAD51^HA-AID^ tachyzoites were incubated for 4 hours with HU and CPT, both tested in *T. gondii* before [31–33]. HU is a ribonucleotide reductase inhibitor that depletes intracellular deoxynucleotide triphosphates, blocking DNA synthesis at the beginning of the S phase, triggering the intra-S phase checkpoint [34, 35]. CPT blocks the rejoining step of the cleavage / religation reaction of topoisomerase I, stabilizing the topo1/DNA complex [36]. This causes the replication forks to collide with each other and with the transcription machinery, affecting the DNA synthesis preferentially at mid and late S-phase [37]. In these assays, the percentage of tachyzoites with TgRAD51 foci as well as number of foci per nucleus were analyzed. Only in the presence of HU were we able to observe a significant increase in the percentage of tachyzoites with foci and in the number of foci per nucleus (Fig. 2B, C, D, S4). These results indicate that TgRAD51 foci are associated with the generation of DNA replication stress in intracellular parasites that are actively going through their cell cycle.

### Lack of TgRAD51 alters tachyzoite replication

To determine the relevance of TgRAD51 on tachyzoite replication, the replication rate of RAD51^HA-AID^ was analyzed in intracellular tachyzoite cultures. In the presence of IAA, we observed a slight decrease in the number of RAD51^HA-AID^ tachyzoites compared to the parental (Fig. 3A). A more pronounced defect when we analyze the number of PV with an abnormal number of tachyzoites. In addition to the typical 1, 2, 4, and 8 tachyzoites per vacuole, a significantly higher number of PV containing 3, 5, 6, or 7 tachyzoites were observed in the presence of IAA compared to the control (Fig. 3B). Using an anti-acetylated tubulin (acTubulin) antibody, which labels daughter cells [38], we were able to observe replicating tachyzoites with only one daughter cell or PV with only one of the two or three tachyzoites entering into replication (Fig. 3C). This would indicate that although TgRAD51 is not essential for tachyzoite replication overall, some tachyzoites experience trouble synchronizing replication.

**Fig. 3.**
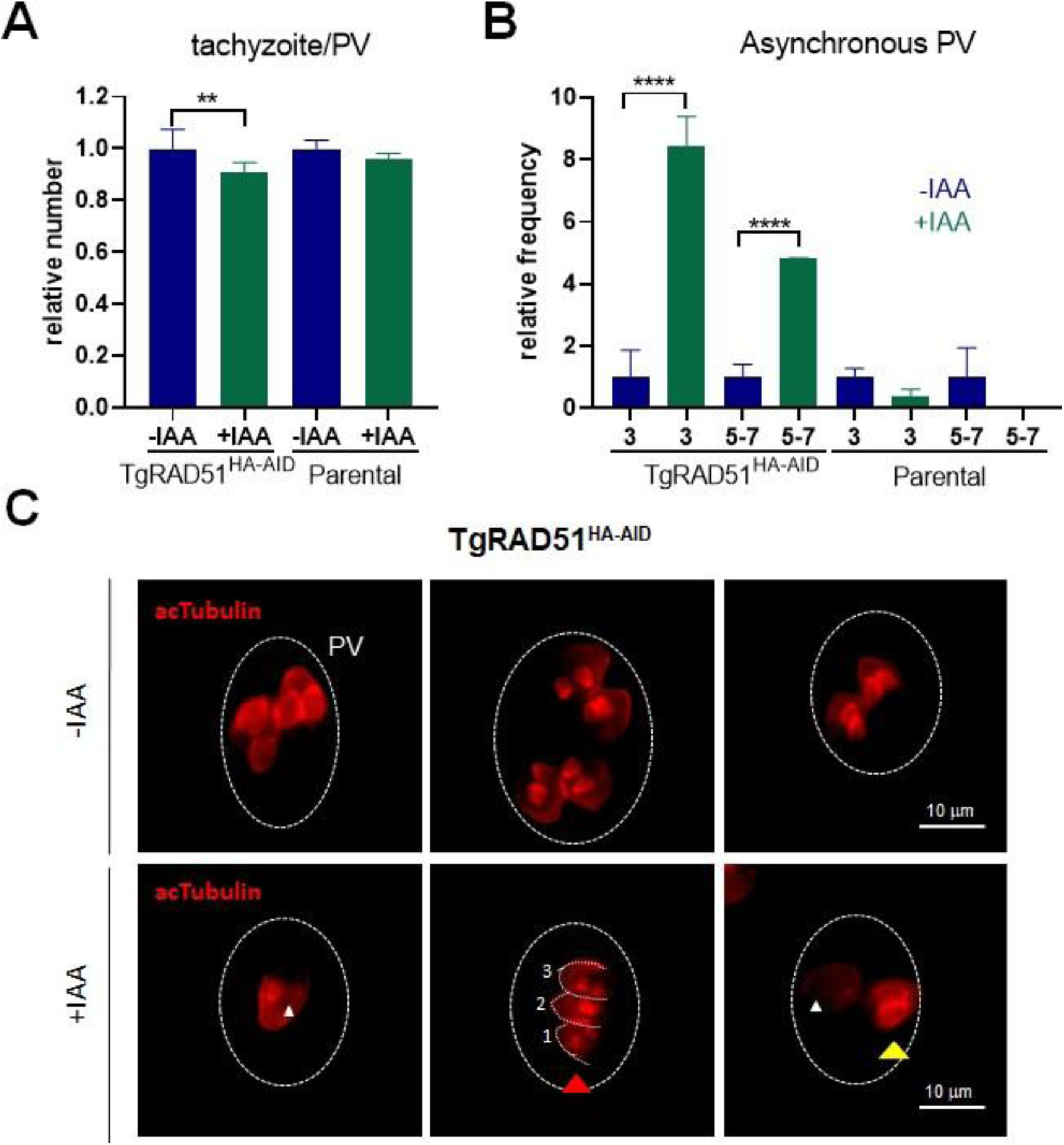
Analysis of *T. gondii* replication in the absence of TgRAD51. hTERT monolayers infected with RAD51^HA-AID^ tachyzoites or the parental strain were treated in the presence of auxin (+IAA) for 16 h, or without treatment (- IAA). The number of tachyzoites per parasitophorous vacuole (PV) was counted in 100 PVs for each case. Each assay was done by triplicate. **A.** The relative average of tachyzoites per PV was analyzed for comparative analysis. The values obtained in the treated (IAA+) and untreated (IAA-) *T. gondii* lines were divided by the average of the untreated ones. The relative average and statistical analysis were done separately for RAD51^HA-AID^ and the Parental strain. The relative data were analyzed by the one-way ANOVA test and Tukey’s multiple comparisons. **p<0.01. The graph is representative of three different assays with similar results. **B.** The frequency of PV with abnormal numbers of tachyzoites that may denote loss of replication synchronization was analyzed. Statistical analysis was done separately for RAD51^HA-AID^ and the Parental strain. Data were analyzed by one-way ANOVA and Tukey’s multiple comparison test. ****p<0.0001. The graph is an example of three assays with similar results. **C.** Intracellular RAD51^HA-AID^ tachyzoites without (-IAA) and with auxin (+IAA) were labeled with anti-acetylated tubulin (acTubulin) antibody to detect daughter cells budding in replicating tachyzoites. White arrowhead: indicates the lack of daughter cell budding within a replicating tachyzoite. Red arrowhead: shows a PV with three tachyzoites. Yellow arrowhead: shows a tachyzoite without daughter cell budding next to another one replicating within the same PV, indicating loss of synchronization.

To test the possibility of arrest in the cell cycle progression, we analyzed *T. gondii* proliferating cell nuclear antigen 1 (TgPCNA1) protein following TgRAD51 knockdown. TgPCNA1 is located in the nucleus but displays a diffuse pattern during the G1 phase and a punctuated pattern in S-phase [11, 39]. The anti-TgPCNA1 antibody was tested on intracellular RAD51^HA-AID^ tachyzoites without IAA, and the correct discrimination of the cell cycle phases could be verified (Fig 4). In the presence of IAA, it was possible to detect tachyzoites in S-phase and others in G1-phase in the same PV, indicating that the synchronicity in replication is lost when TgRAD51 is diminished, with G1 arrest for some tachyzoites.

**Fig. 4.**
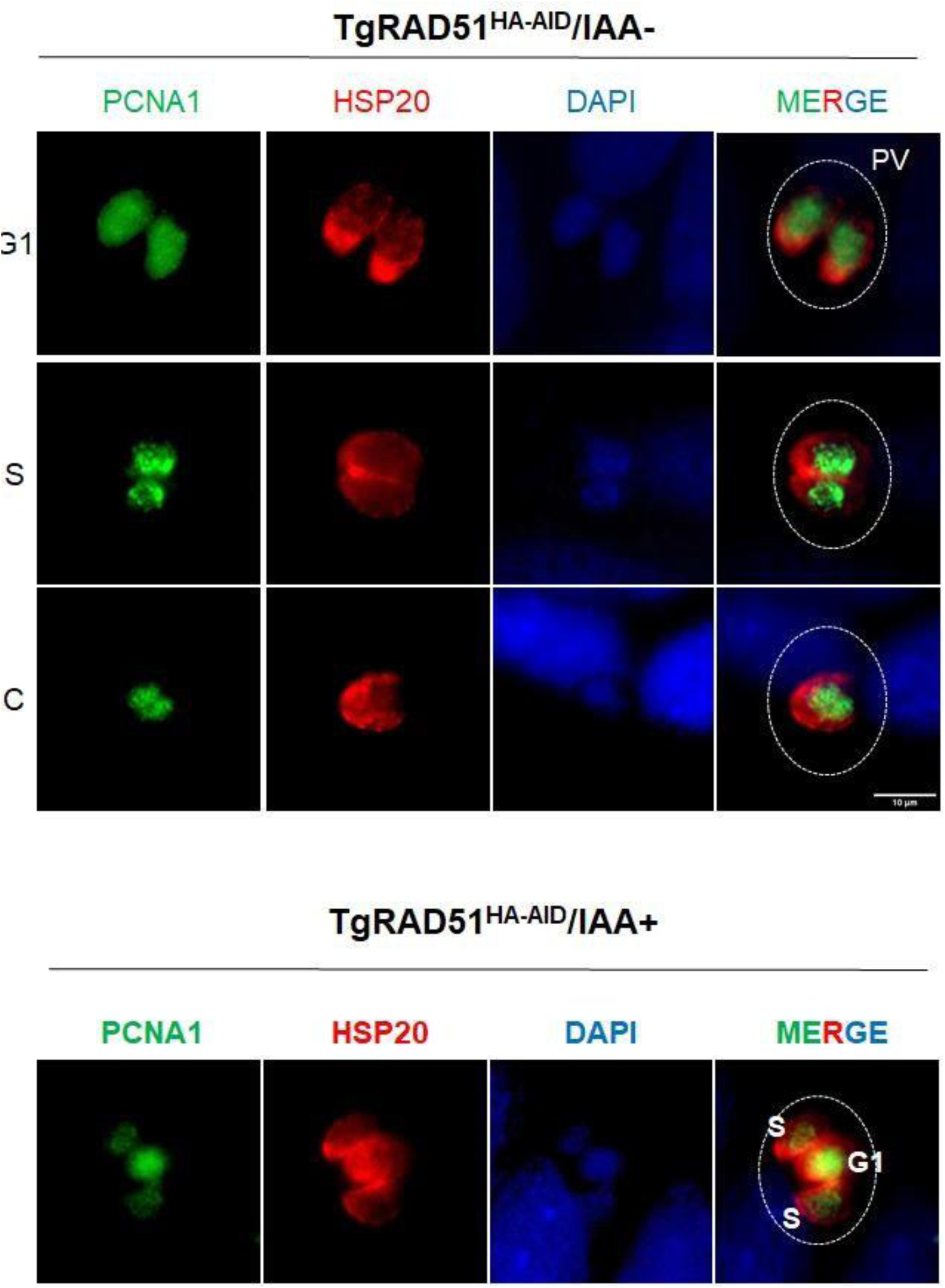
Cell cycle analysis of RAD51^HA-AID^ in the absence of TgRAD51. Intracellular tachyzoites were incubated overnight without or with IAA. The cell cycle phase of the tachyzoites was analyzed by labeling with PCNA1. Thus, it is possible to see in the control without auxin (IAA-) that PCNA1 can distinguish between G1 (diffuse label in the nucleus), S (PCNA1 foci), and cytokinesis (C, PCNA1 foci and shrunken nucleus). In IAA-culture, tachyzoites from each PV replicate synchronously, but not between PVs. In the presence of IAA, PVs with tachyzoites in different phases of the cell cycle are observed, indicating a loss of synchronization due to delay or arrest of the cell cycle. The white bar indicates 10 μm. This bar is the same for all panels.

### RAD51 inhibitor B02 alters *T. gondii* replication

Based on protein 3D modeling, Vydyam et al. [24] identified a compound called B02 as an anti-PfRAD51 inhibitor with an IC_50_ of 3-8 μM. Notably, the plasmocidal effect of B02 occurs at a lower concentration than against mammalian cells. B02 did not show activity against *Entamoeba histolytica* EhRAD51 [40]. Due to these differences in RAD51 sensitivity to B02 between species, we performed a sequence similarity analysis between the different RAD51s. All RAD51s are highly conserved, mainly from the HHH domain to the C-terminus (Fig. 5, Fig S5). However, TgRAD51 and PfRAD51 exhibit greater similarity to each other compared to HuRAD51, whereas EhRAD51 is the most divergent homolog examined (Fig. 5A). We reasoned that B02 could be a good candidate against TgRAD51.

**Fig. 5.**
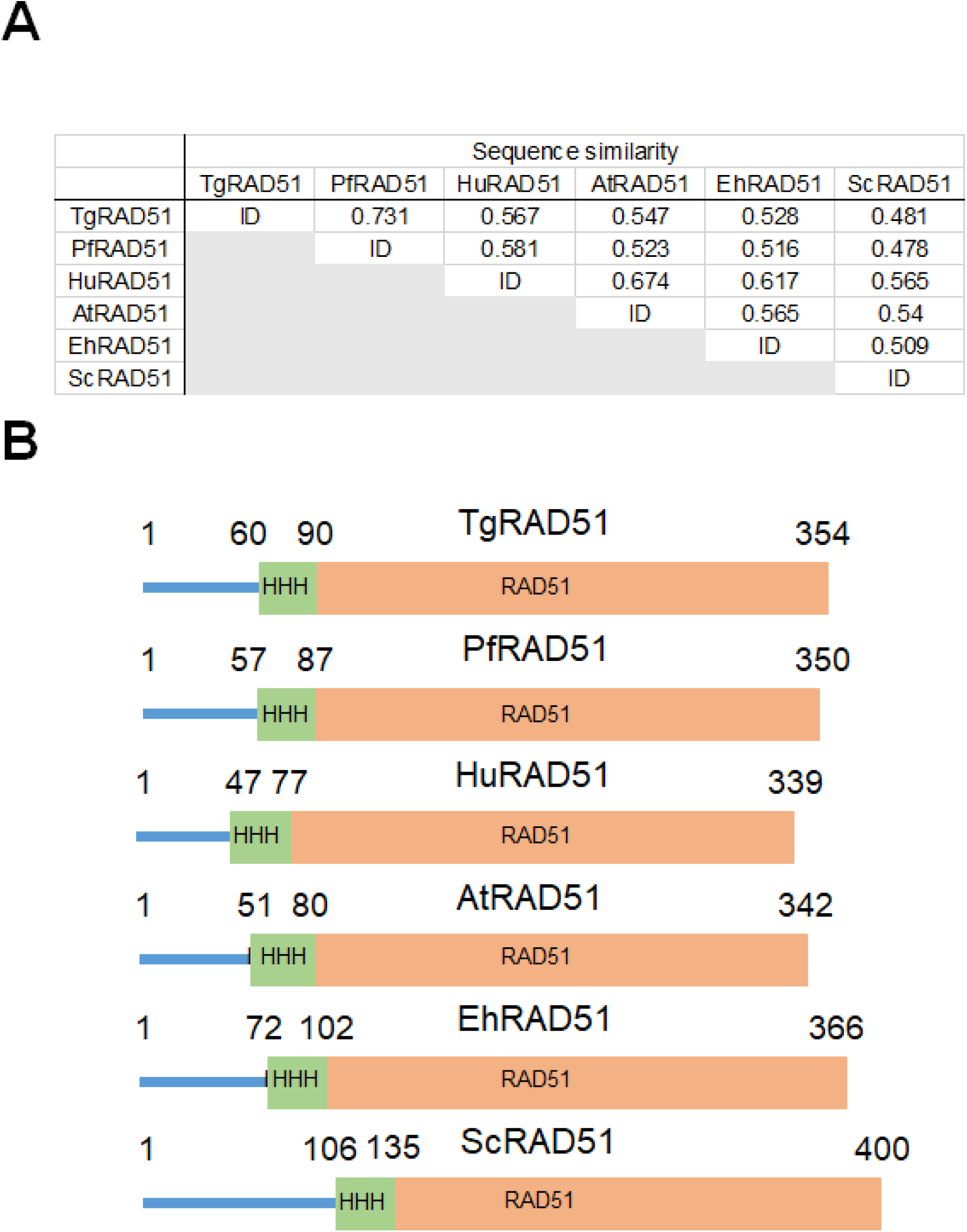
Similarity of TgRAD51 with RAD51 from other species and B02 response. **A.** Sequence identity matrix analyzed in Bioedit version 7.2.5. Sequence alignment was performed using Clustal W multiple alignment, boostrap NJ tree: 1000. Identities were calculated using Sequence Identity Matrix. TgRAD51 (ToxoDB geneID TGME49_272900), *Plasmodium falciparum* RAD51 (PfRAD51, Plasmodb GeneID PF3D7_1107400), *Entamoeba histolytica* RAD51 (EhRAD51, GenBank ID XM_648984), Human RAD51 (HuRAD51, uniprot Q06609), *Arabidopsis thaliana* RAD51 (AtRAD51, uniprot P94102) and *Saccharomyces cerevisiae* RAD51 (ScRAD51, uniprot P25454). **B.** TgRAD51 shares structural and sequence length similarity with *Pf* RAD51, EhRAD51, HuRAD51, AtRAD51 and ScRAD51. Motif and domains were analyzed by Motifscan web page (https://myhits.sib.swiss/cgi-bin/motif_scan): HHH, Helix-hairpin-helix motif; RAD51: RAD51 domain.

Concentrations of B02 lower than 15 μM have no obvious cytotoxic effect on hTERT host cells which showed a CC_50_ ∼50 μM (Fig. 6A). To test the anti-*T. gondii* effect of B02, intracellular RH-RFP, a strain in which both HRR and NHEJ pathways are present, were incubated for 72 hours in the presence of increasing drug concentrations (0 to 15 μM). A dose-dependent inhibition of *T. gondii* growth was detected with an IC_50_ of 4.8 μM and a selectivity index (SI) of ∼10.4 (Fig. 6B). The IC_100_ value is reached at a concentration of 15 μM, below the CC_50_ value.

**Fig. 6.**
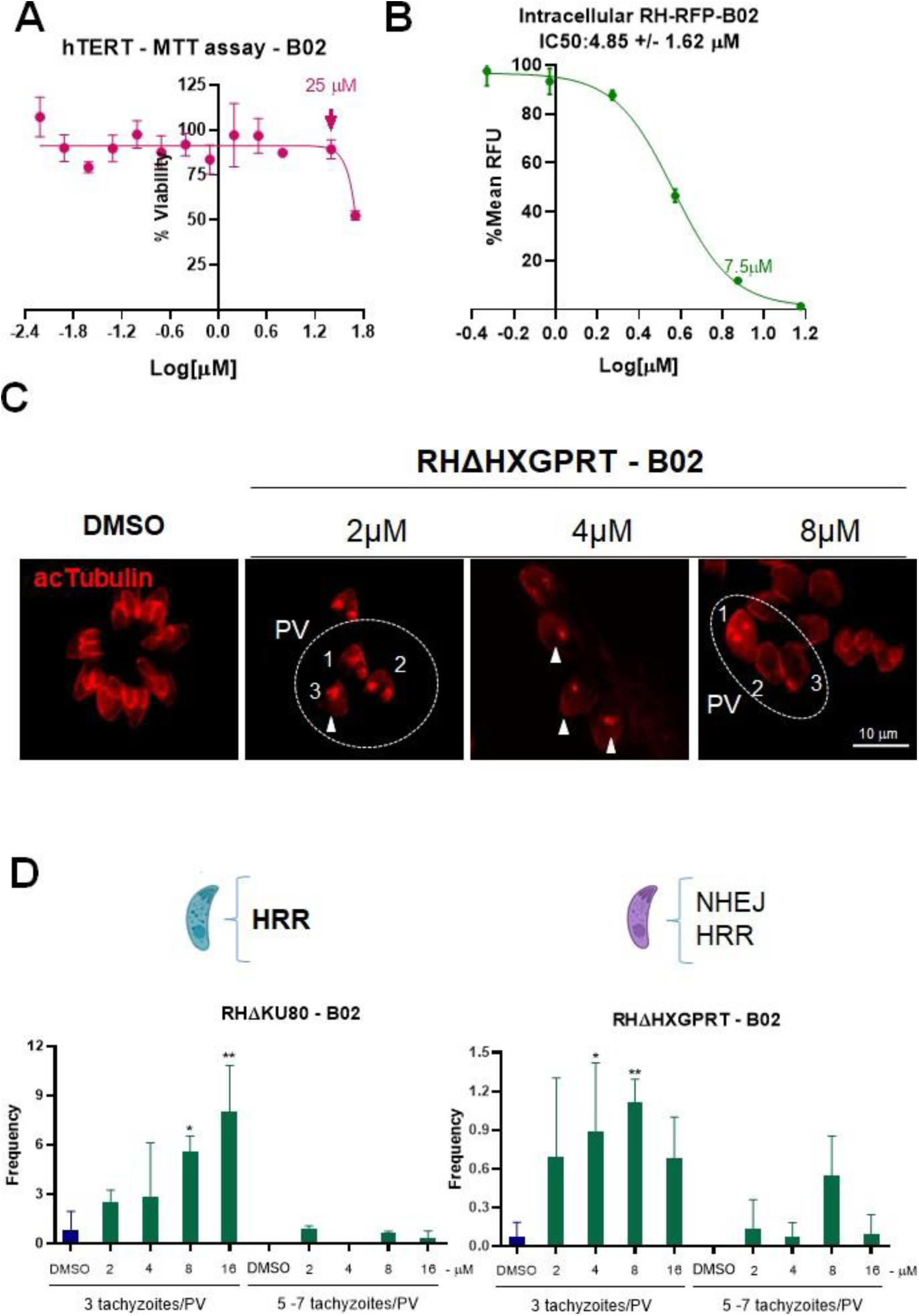
Effect of RAD51 inhibitor B02 on *T. gondii* growth. **A.** Cytotoxic concentration 50% (CC_50_) was calculated on hTERT monolayers grown in increasing concentrations of B02 for 72 h. The MTT reagent (3-(4,5-dimethylthiazol-2-yl)-2,5-diphenyl-2H-tetrazolium bromide) was used to analyze the metabolic status of hTERT cells and determine cell viability. The percentage (%) viability was graphed. CC_50_ value was calculated with GraphPad Prism 8 (log[inhibitor] vs. response -- Variable slope. least squares fit). A CC_50_ ∼50μM was obtained based on two independent assays. As a reference, 25 μM concentration is indicated. **B.** The half-maximal inhibitory concentration (IC_50_) was calculated on hTERT monolayers infected with *T. gondii* RH-RFP strain grown in increasing concentrations of B02 for 72 h. Analysis was done by measuring RFU at excitation 544 nm; emission 590 nm. IC_50_ was calculated with GraphPad Prism 8 (log[inhibitor] vs. response -- Variable slope. least squares fit). An IC_50_ of 4.85 μM was obtained based on three independent assays. As a reference, 7.5 μM concentration is indicated. **C.** hTERT monolayers infected with RHΔHXGPRT tachyzoites treated in the presence of B02 or with solvent alone (DMSO) were labeled with anti-acetylated Tubulin (acTubulin) antibody to detect daughter cell budding in replicating tachyzoites. The number of tachyzoites within a PV is indicated inside the panel. The white arrowhead at 2 μM panel shows the lack of daughter cell budding within a replicating tachyzoite. The white arrowhead at 4 μM panel shows the lack of daughter cell budding within a replicating tachyzoite. At 8 μM panel: 1. one replicating tachyzoite with three daughter cell buddings is shown. 2 and 3: tachyzoites without daughter cell budding in a PV with another replicating tachyzoite, indicating loss of synchronization or arrest. **D.** The frequency of PVs with abnormal numbers of tachyzoites that may denote loss of synchronization in replication was analyzed under B02 treatment. Statistical analysis was done for the strains RHΔHXGPRTΔku80 (RHΔKu80) and RHΔHXGPRT. Data were analyzed by one-way ANOVA and Tukey’s multiple comparison test. *p<0.05; **p<0.01. The graph is an example of three independent experiments with similar results. HRR: homologous recombination, NHEJ: non-homologous end joining. As shown, in one of the strains (purple) both DSB repair pathways are available, while in the other (blue) only the HRR pathway is present.

We next analyzed the effects of B02 treatment on parasites RHΔKU80 and RHΔHXGPRT by IFA using anti-acTubulin antibody. In the presence of B02, vacuoles containing 3 tachyzoites could be observed, in some cases with one daughter cell budding, as well as a loss of synchrony in replication (Fig. 6C). In RH strain, the frequency of PVs with abnormal numbers of tachyzoites increased in a dose-dependent manner, with most PVs housing 3 tachyzoites (Fig. 6D). This replication abnormality after B02 treatment was also observed in RHΔKu80, with an even higher number of 3 Tz/PV. These results show that B02 affects tachyzoites at a lower concentration than host cells, causing anomalous parasite replication.

### B02 arrests *T. gondii* cell cycle at S-phase

To further examine this replication defect in response to TgRAD51 inhibition, RH tachyzoites were grown in 5 μM B02 for 6 h and cell cycle stages were determined by FACS analysis. We observed a significant arrest in S phase during B02 treatment (Fig. 7A). As expected, treatment with the positive control hydroxyurea (HU) arrested parasites in G1. We noted that B02 generated a shoulder in the cytometry profile near the M-phase peak, compatible with the end of S-phase (Fig. 7B). To study this peak, we divided the S phase into two: S-early phase starting at G1 up to the middle of S-phase and S-late phase starting from the middle of S-phase to mitosis. We analyzed the frequency of both sub-cycles, finding that B02 induced arrest at S-late (Fig. 7C). This suggests that B02 largely affects *T. gondii* when the largest amount of DNA has been replicated, which is necessary to activate the HHR pathway.

**Fig. 7.**
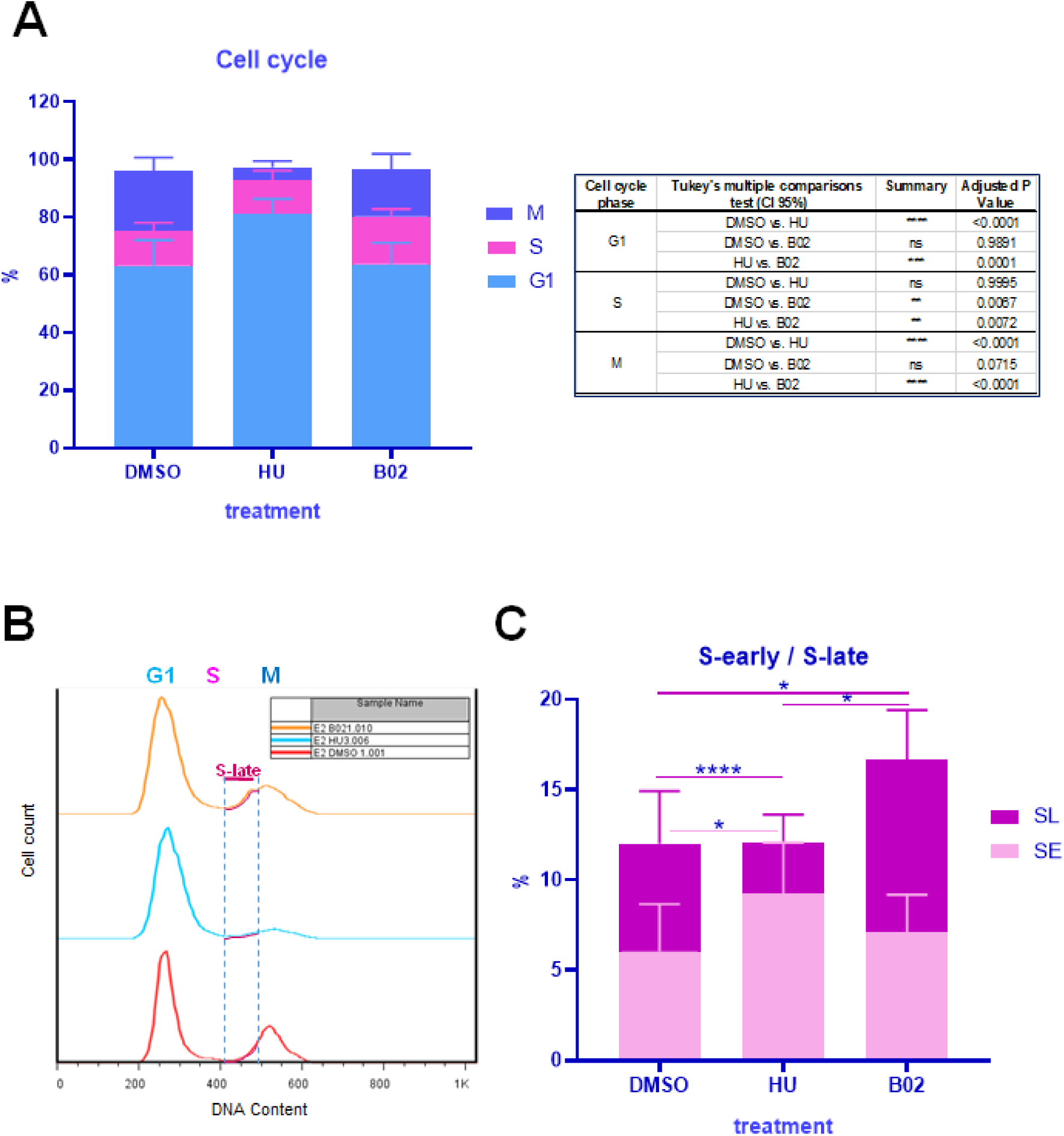
Analysis of RHΔHXGPRT tachyzoites cell cycle after treatment. A. hTERT monolayers infected with RHΔHXGPRT tachyzoites were incubated with 5 μM B02, 4 mM HU and DMSO 0.1% for 6 h. Propidium iodide was used to stain DNA. DNA content of tachyzoites were analyzed by FACS to determine the G1 (1N), S (>1-1.8N) and M (2N) cell cycle phases. The number of tachyzoites in each phase was determined and their percentage was plotted. Data from three independent experiments were used. Statistical analysis was performed with one-way ANOVA and Tukey’s multiple comparison test (see table at right). **B.** Number of tachyzoites along the cell cycle were plotted. The graph is an example of three independent experiments with similar results. The S-late region is shown. **C.** Bar graphs showing the percentage (%) of tachyzoites in each S-phase of the cell cycle. Statistical analysis was performed with one-way ANOVA and Tukey’s multiple comparison test (****p < 0.0001, *p < 0.05).

### B02 induced bradyzoite differentiation

Different types of stress induce tachyzoite to bradyzoite differentiation *in vitro* [41, 42]. We therefore tested whether the RAD51 inhibitor B02 prompts differentiation as a result of DNA replication stress. We treated tachyzoites of the cystogenic strain Me49 with B02 for 96 hours under normal culture conditions. Me49 tachyzoites treated with DMSO and grown under normal conditions were used as negative control (C-). As positive control (C+), we induced bradyzoite differentiation by alkaline stress (pH 8.1). The differentiation process was quantified by labeling cyst wall with *Dolichos biflorus*-*agglutinin* (DBL+). In the presence of different doses of B02 (5 and 10 μM), a significant positive DBL labeling was observed in comparison to C-, and similar to that observed in the C+ (Fig. 8A).

**Fig. 8.**
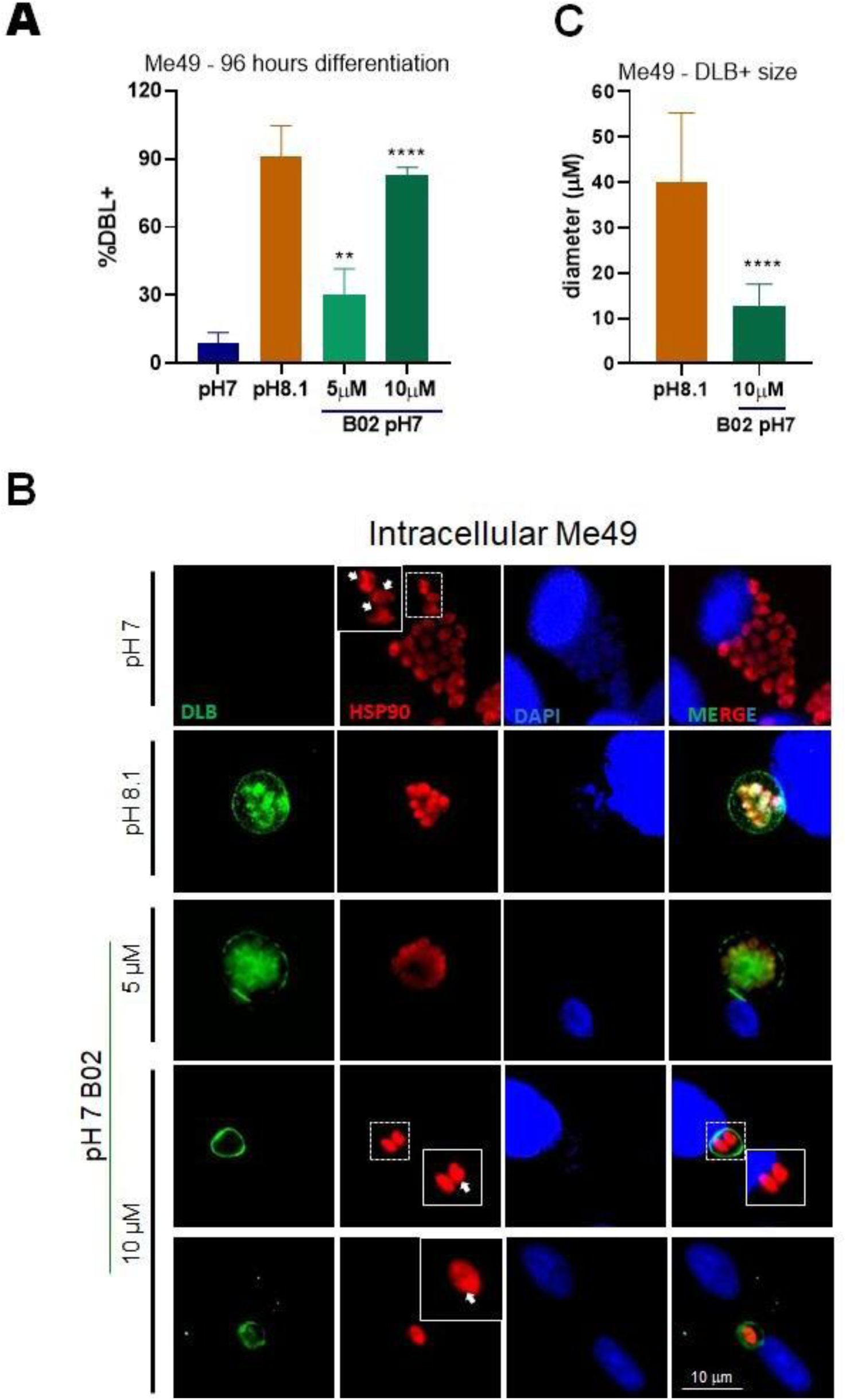
Bradyzoite induction analysis in B02-treated *T. gondii*. Monolayers of hTERT infected with tachyzoites of the cystogenic strain Me49 were treated with different doses of B02. *T. gondii* treated with dissolvent (DMSO) were used as a negative (basal) control (C-). Intracellular tachyzoites were incubated for 96 hours under normal culture conditions (37°C, 5% CO2, pH 7). As a positive control (C+) a group of infected host cells were incubated at pH 8.1 to induce differentiation by alkaline stress. At the end of the experiment the infected cell cultures were analyzed by epifluorescence microscopy using the cyst wall marker *Dolichos biflorus*-agglutinin (DLB) and anti-TgHsp90 antibody (cytosolic in the tachyzoite, cytosolic and nuclear in the bradyzoite). **A.** The number of cyst-like structures (DLB+) was counted by triplicate in 100 PVs taken at random. Statistical analysis was performed with one-way ANOVA and Tukey’s multiple comparison test. Only B02 vs C-was analyzed. (****p < 0.0001, **p < 0.01). This is an example of three independent experiments with similar results. **B.** Infected cell cultures were analyzed by epifluorescence microscopy. In the treatment with 10 μM B02, an image of a DBL+ structure with 1 bradyzoite and another with 2 bradyzoites are shown. The white arrow shows a zoom of tachyzoites from a PV for TgHSP90 labeling. At pH 7, the label is observed outside the tachyzoite nucleus. In conditions inducing differentiation (pH 8.1 and B02) the TgHSP90 label covers the entire bradyzoite. **C.** The diameter of 25 DBL+ structures was analyzed for 10 μM B02 and pH 8.1 (C+) for each treatment in triplicate and graphed. Statistical analysis was performed with the Student T test. ****p < 0.0001.

To further examine bradyzoite conversion, we monitored TgHsp90 localization in response to B02 treatment. TgHsp90 is cytosolic in tachyzoites but cytosolic and nuclear in bradyzoites [43]. The subcellular localization of TgHsp90 staining in Me49 at pH 8.1 or treated with B02 is compatible with that expected for the bradyzoite, even in DBL+ structures with 1 or 2 *T. gondii* cells (Fig. 8B), further suggesting initiation of differentiation in these parasites.

Remarkably, in the presence of 10 μM B02, small cyst (DBL+) structures containing only 1 or 2 *T. gondii* cells were observed (Fig. 8B). Of note, since at this concentration most *T. gondii* die, the number of PV/cyst-like structures is much lower than with 5 μM B02 or the DMSO control. Cyst size counting in the different treatments and controls showed that as the B02 dose increases, cyst size decreases, being significantly smaller in 10 μM B02 (Fig. 8C). We decided to analyze the presence of the tachyzoite marker SAG1 in the DLB+ vacuoles as an indicator, when absent or less intense (SAG1-), of mature cysts (Fig. 9). In the presence of 5 μM B02 41,7% of cyst-like structures were compatible with mature cyst (DBL+/SAG1-). In the 10 μM treatment, with smaller DLB+ vacuoles, only 8,8% were DBL+/SAG1-. These data suggest that B02 can produce mature cysts.

**Fig. 9.**
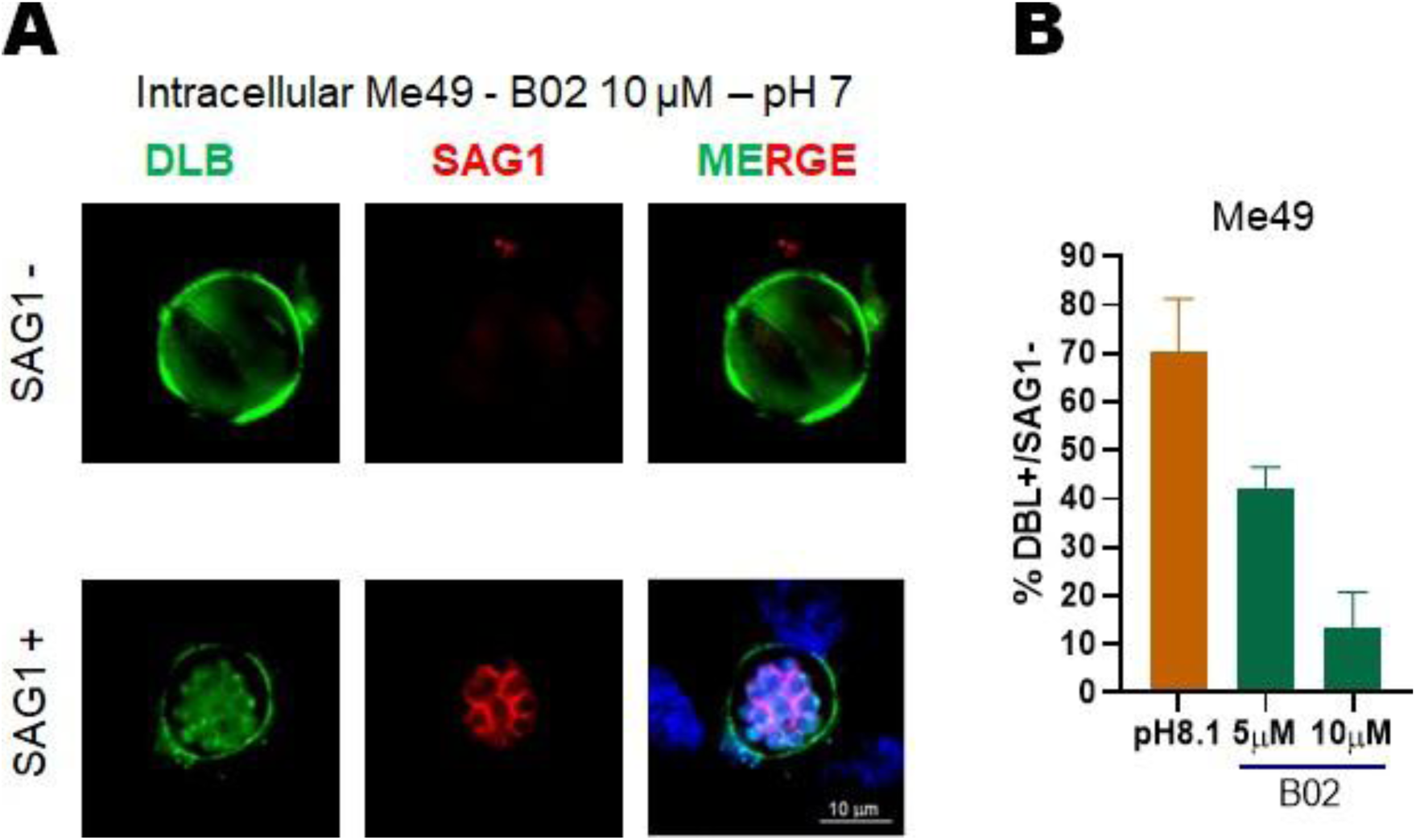
Analysis of mature cyst formation in B02-treated *T. gondii*. SAG1 is a tachyzoite-specific protein which decreases in the mature bradyzoite until it becomes null. Therefore, its detection in DLB+ structures (DLB+/SAG1+) would be indicative of an immature cyst, while its slight or null detection (DLB+/SAG1-) would be indicative of a mature cyst. **A.** Monolayers of hTERT infected with Me49 strain tachyzoites were treated with 10 μM B02 for 96 h. They were subsequently analyzed by epifluorescence microscopy. DBL was used to label cyst structures and anti-SAG1 antibody was used as tachyzoite marker. SAG1-would correspond to mature cysts. **B.** Me49 tachyzoites induced to differentiate at pH 8.1 or BO2 were labeled with DLB+ and the anti-SAG1 antibody. In each case, at least 50 DLB+ structures were randomly selected and analyzed for the presence of SAG1, in fields where tachyzoites were present so as to compare the intensities.

### Effect of CPT or HU and B02 combinations against *T. gondii* growth

As mentioned, CPT is a genotoxic drug that affects the growth of *T. gondii* with an IC_50_ of 5 μM [32]. Thus, incubation of intracellular tachyzoites with CPT should generate DNA damage that leads to increased DNA replication stress, DSB formation, and HRR activation. In order to test whether the combination of CPT with B02 improved the anti-*T. gondii* effect, RH RFP tachyzoites were incubated with 2.5 μM B02 and CPT (Fig. 10). As can be observed, the combination of CPT and B02 significantly improved the anti-*T. gondii* effect compared to the use of the drugs administered individually.

**Fig. 10.**
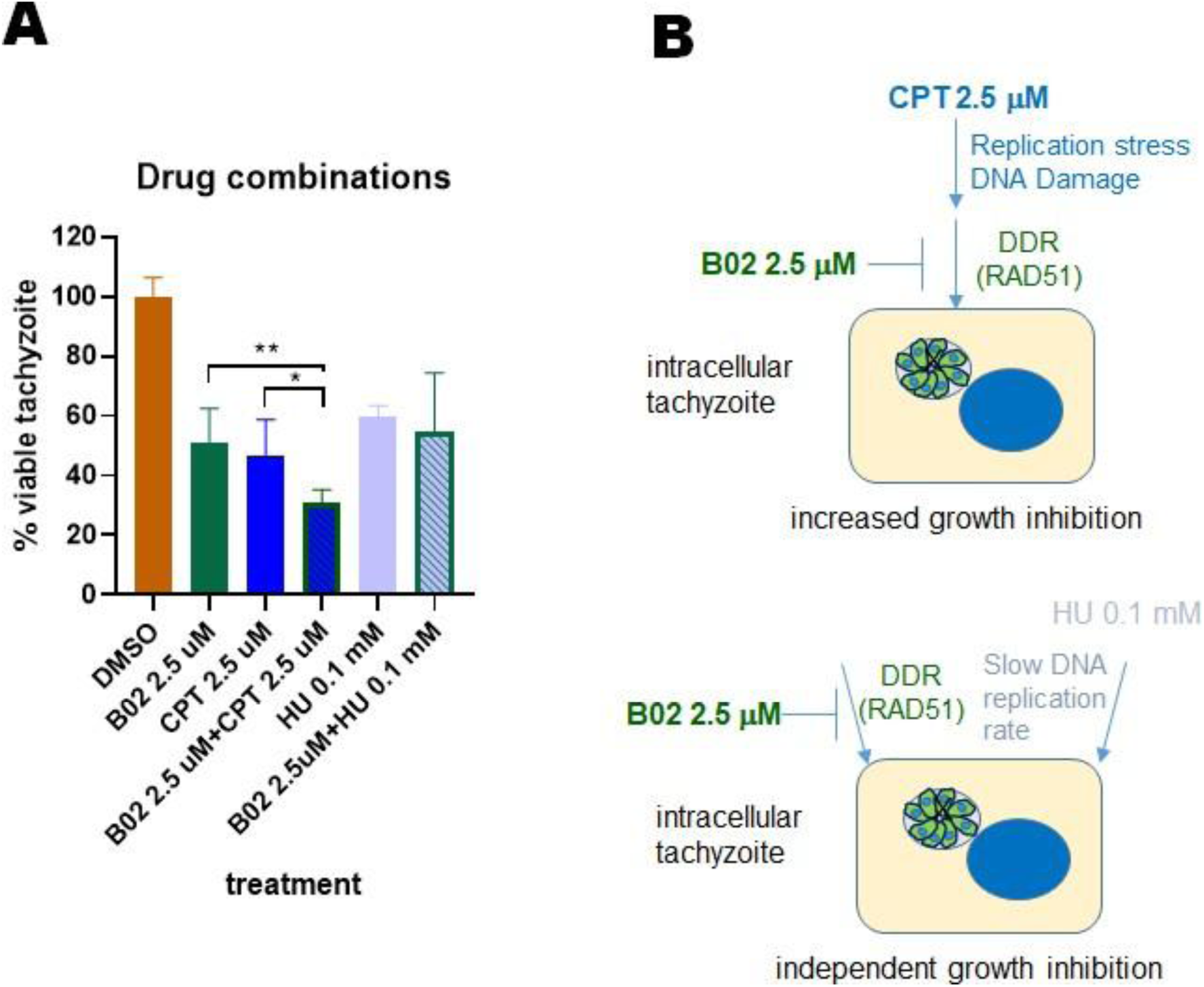
Effect of the combination of CPT and B02 on *T. gondii* growth. **A.**. Intracellular RH RFP tachyzoites were incubated for 96 hours in the presence of 2.5 μM B02, 2.5 μM CPT, 0.1 mM HU and combinations thereof. Analysis was done by measuring RFU at excitation 544 nm; emission 590 nm. Statistical analysis was performed with one-way ANOVA and Tukey’s multiple comparison test. *p < 0.05, **p < 0.01. The graph is an example of three independent assays with similar results. **B.** Above: CPT generates DNA replication stress, leading to DSBs that must be repaired mainly by HRR. By inhibiting RAD51 with B02, altering the DNA Damage Response (DDR), the repair process cannot overcome the damage, thereby increasing the anti-*T gondii* effect. Below: While B02 inhibits RAD51, blocking the DDR, HU at this low concentration would be generating an unknown anti-*T. gondii* effect, but not associated with blocking the cell cycle. For this reason, some tachyzoites are eliminated or blocked in one metabolic pathway and others in another, without the enhanced effect of the drugs when combined.

In a similar manner, we analyzed the effect of the combination of HU with 2.5 μM B02. Since high doses of HU generated an inhibitory effect with high cell death in cultures grown for 96 hours, a concentration of 0.1 μM was used. In this case, we did not observe differences in the growth of the parasites with respect to their drugs administered individually (Fig. 10). This indicates that the drugs would not intervene in the same stage of the cell cycle, either generating damage or affecting its repair.

## Discussion

In this study, we used genetic and pharmacological approaches targeting RAD51 to investigate HRR in *T. gondii*. Since RAD51 is a surrogate marker of HRR functionality, we generated a *T. gondii* line RAD51^HA-AID^ where RAD51 protein declines in the presence of auxin. The phenotypic analysis of RAD51^HA-AID^ showed that this protein is important for correct tachyzoite replication under normal growth conditions. This result was validated in studies using the RAD51 inhibitor B02. We conclude that TgRAD51 inhibition can lead to replicative stress that arrests cell cycle in late S phase, possibly initiating bradyzoite differentiation as an adaptive response. These results indicate a role for TgRAD51, likely through HRR pathway, in the normal growth of tachyzoites.

We observed RAD51 foci in intracellular tachyzoites grown under normal conditions without genotoxic agents. Similar findings are observed in other eukaryote systems where it is possible to identify RAD51 foci in immortal cells grown without genotoxic agents [44]. However, no nuclear RAD51 foci were observed in trophozoites of *E. histolytica* collected at the exponential growth phase [45]. It is known that in highly replicating cells, such as tumor cells, DNA replication stress is generated that can lead to fork collapse and DSB generation triggering the HRR response [46]. We therefore consider that the RAD51 foci in *T. gondii* are due to the high replication rate of tachyzoites [11]. The absence of foci in extracellular non-replicative tachyzoites, as well as the increment observed in the presence of HU, also support this idea.

It is interesting that only HU increased the number of tachyzoites with TgRAD51 foci as well as the number of foci per nucleus under the tested conditions. HU clearly shows to arrest parasites in G1, whereas CPT was described to arrest cells in G2/M [47]. Moreover, HU activates the S-phase CHK1 checkpoints slowing down replication [48]. CPT would be associated with activation of the late S-phase checkpoint [37, 49]. This could explain why under the conditions of the assay, with tachyzoites incubated for 4 hours with drugs, TgRAD51 foci increase was only observed in tachyzoites treated with HU and not with CPT.

In addition to its role during HRR, RAD51 has other non-canonical roles [50]. Among them, RAD51 is required for the generation of fork reversal mechanism [51], facilitates post-replicative repair by translesion [52], and promotes RNA:DNA hybrid (RNA loop) formation [53]. RAD51 can bind to Telomeric Repeat-containing RNA (TERRA), favoring its binding to the telomeric regions of chromosomes [54]. Of note, preliminary laboratory studies indicate that *T. gondii* subtelomeres may be expressing TERRA lncRNA (unpublished data). With a knockdown now in hand, future studies can address whether TgRAD51 exhibits any of these non-canonical functions in *T. gondii*.

Knockdown of TgRAD51 leads to replication defects including a reduction in replication rate, loss of synchronization, and cell cycle arrest. Similar effects have been reported in *Leishmania major* and *Schizosaccharomyces pombe* [55, 56]. The lack of LmRAD51 and SpRAD51 did not affect cell viability but slowed growth. In mice, the lack of RAD51 affects embryogenesis, with a probable role in cell proliferation [57]. These findings further support the relevance of RAD51 in the normal DNA replication process of diverse organisms, and the identification of RAD51 orthologues in early-branching eukaryotes suggest an ancient development of robust DNA repair mechanisms in cellular evolution.

Consistent with data obtained after genetic manipulation, the RAD51 inhibitor B02 produced a similar negative effect on *T. gondii*. B02 induced replication alterations that generated PVs with abnormal numbers of tachyzoites. B02 caused an arrest in the S-late phase in the lytic cycle of tachyzoite. This is expected since the drug inhibits RAD51, an enzyme that must repair DNA damage via the HRR pathway, which occurs in S-late [58, 59]. Knockout of the LmRAD51 gene showed that LmRAD51 is associated with S-phase entering G2/M [56]. B02 also drastically reduced the number of tachyzoites per PV. Complete growth inhibition of *T. gondii* was achieved at doses of B02 (15 μm) that are below the cytotoxic effect on host cells. Furthermore, the IC_50_ of B02 against *T. gondii* was 4.8 μM, similar to the IC_50_ observed for *P. falciparum* (3 - 8 μM), and quite far from the IC_50_ (27.4 μM) observed for human RAD51 [22, 24]. These findings suggest that B02 is selectively toxic against apicomplexan RAD51 relative to human RAD51, elevating it as an attractive anti-parasitic compound.

A relevant effect of B02 is the induction of mature cyst-like structure in intracellular tachyzoites. It is known that in vitro different stress situations induce the conversion of tachyzoite to bradyzoite [60, 61]. All stress mechanisms that promote differentiation seem to be a response of *T. gondii* to an environmental stress that may globally affect different metabolic pathways [62]. However, in this case the mechanism would be associated with DNA replication stress, indicating that *T. gondii* is capable of finely sensing its cell cycle, and any modification of the replicative process could trigger a differentiation response. So far, the current model is that although the differentiation process could start in the G1 phase, the bradyzoite form appears at the end of the S phase to enter into an arrest in G0 [63]. Thus, it is understood that at least 2 bradyzoites should appear from an induced tachyzoite during differentiation. Moreover, in parasites arrested in G1, either using pyrrolidine dithiocarbamate or in RH^TK+^ line treated with thymidine, differentiation to bradyzoite is blocked [64]. In this sense, the presence of some cysts containing only one bradyzoite in presence of B02 would be in contradiction with that model. Therefore, we consider that tachyzoites are likely attempting to differentiate into bradyzoites, but arrest in the S phase prevents the completion of the process.

The combination of CPT with B02 increases the inhibitory effect of the treatment on *T. gondii* growth. The use of drug combinations with B02 is not new [65], but the genotoxic agent CPT had not been used. CPT is not FDA-approved but other CPT derivatives, irinotecan and topotecan, are used for human treatments [49, 66]. So far, only topotecan was tested as an anti-*T. gondii* drug showing effects at high doses [67]. The use of similar genotoxic agents more specific for *T. gondii*, which are not toxic to the host cell, could be an alternative to combine with B02 in future trials of new anti-*T. gondii* therapies.

In contrast, the combination of 0.1 mM HU and 2.5 μM B02 does not improve the anti-*T. gondii* effect. This may be due to the very low dose of HU used. The effect of HU on cell proliferation varies greatly depending on the dose. In other cell types, HU causes G1 arrest at concentrations higher than 200 mM [68]. At these concentrations, reactive oxygen species (ROS) are also generated, which can cause oxidative DNA damage. At lower concentrations, it may have other effects, such as altering the Fe-S centers of several cellular enzymes [48]. At concentrations of 0.5 mM, HU can induce telomere dysfunction [69]. All these data indicate that perhaps the effect of HU at 0.1 mM is different from that generated with 1 mM. In *T. gondii*, HU is a potent inhibitor of *T. gondii* growth even at concentrations much lower than those used in mammals, as observed in our experiments. In the past, Kasper and Pfefferkorn [70] had already demonstrated the potent anti-*T. gondii* effect, determining that 18 mg/ml (0.234 mM) of HU completely blocks the growth of *T. gondii*. However, this concentration did not severely affect DNA synthesis, although a greater decrease was observed at higher concentrations. In conclusion, it is possible that the concentration of 0.1 mM does not generate damage associated with DNA replication, and for that reason an improvement in the anti-*T. gondii* effect could not be observed when HU was combined with B02.

## Conclusion

We observed that TgRAD51 plays a relevant role in replicating tachyzoites under normal culture conditions without exogenous DNA damaging agents being added. Both the lack of TgRAD51 and its inhibition with B02 alter the parasite’s replication rate, cell cycle, and formation of daughter cells. Our data, together with previous observations [67], indicate that TgRAD51 facilitates the recovery of tachyzoites from DNA replication stress accrued during the lytic cycle. DNA replication stress appears to induce cyst-like structures. Together, our findings support the idea that TgRAD51 is an attractive target for future drug development, which would improve in combination with a genotoxic agent that acts on DNA replication.

## Materials and Methods

### Parasite culture

Parasites of the RHΔHXGPRT [71], RHΔKU80 [30], RHΔKU80TIR1 [28, 29], RH RFP, which express red fluorescent protein [72], or Me49 strain (kindly provided by Valentina Martin, Escuela de Ciencia y Tecnología, UNSAM, and analyzed by genotyping), were cultured *in vitro*. Human immortalized fibroblasts (hTERT) (BD Biosciences) monolayers [73] were infected with tachyzoites and incubated in Dulbecco’s modified Eagle medium (DMEM, GIBCO) supplemented with 1% fetal bovine serum, penicillin (100 I/ml; GIBCO), and streptomycin (100µg/ml; GIBCO) at 37°C and 5% CO_2_.

### Generation of parasites expressing RAD51 endogenously tagged with HA-AID

To generate the endogenous tagging of RAD51 with the AID system, we followed the protocol previously described (www.bio-protocol.org/e2728). Briefly, the RAD51 genomic sequence (TGME49_272900) was tagged with AID and three C-terminal HA epitope tags in RHΔKU80TIR1 parasites using CRISPR/Cas9-based and double homologous recombination to generate the *T. gondii* line RAD51^HA-AID^. A single guide RNA (sgRNA) sequence near the stop codon of RAD51 was chosen using the EuPaGDT tool (http://grna.ctegd.uga.edu/) and the sequence was cloned into a pCas9-GFP plasmid [74] by site directed mutagenesis using PCR, designated as pCas9-RAD51. *T. gondii* tachyzoites were co-transfected with pCas9-RAD51 along with a PCR amplicon repair template using a Lonza Biosciences Nucleofector. To amplify this donor we used the following pair of primers: forward: CGCCATCGGCGAAGGAGGCATCGGCGACTACGAAGACAAccgctagcaagggctcggg (P1, forward), and: ctgcatccgtgtagctctgtgactttgagcctgttggaacaaaagctggagctccac (P2, reverse). The repair template encoded a 3xHA-AID tag sequence and a dihydrofolate reductase-thymidylate synthase (DHFR*-TS) selectable marker flanked by 40 bp homologous to either site of the CAS9 cut site to facilitate homology-directed repair of the CAS9-induced break. *T. gondii* parasites were selected for DHFR*-TS integration by three passages in a medium containing 1 mM pyrimethamine (Sigma, SML3579) and cloned by limiting dilution. The modified locus of the transgenic clone was validated by PCR analyses of both 5’ -end and 3’ -end. The primers sequence for validation included the donor primers and an extra pair: gaacatcctcaacaaggaac (P3, forward), and: gccaccgcttgatttttggca (P4, reverse). The validation PCR was executed as explained in Figure S1.

### Immunofluorescence assay (IFA)

Slides with intracellular tachyzoites exposed to the different treatments specified accordingly were grown for 24 h in hTERT on glass slides. All further incubations were carried out at room temperature. Cells were washed in PBS and fixed in 4% paraformaldehyde (PAF) for 20 min. After three washes, cells were permeabilized with 0.2% TX-100 in PBS for 10 min, blocked with 10% PFBS (fetal bovine serum in PBS) for 20 min, then incubated overnight or at 1 h at room temperature with primary antibodies generated at our lab such as: TgHSP90 rabbit (1:500) [43] and TgHSP20 rabbit (1:500) [75]. TgPCNA1 mouse (1:500) was prepared in our laboratory from recombinant *T. gondii* PCNA1 protein expressed in *E. coli*. Commercial antibodies were used as datasheet indicated: Acetylated tubulin (acTubulin) mouse (1:500) from Invitrogen (MA5-18268) and hemagglutinin mouse (anti-HA) (1:200) from Abcam (ab1424). Subsequently, slides were washed several times with PBS and the secondary antibody Alexa Fluor Goat anti-mouse 488 (Invitrogen, 1:2,000) or Alexa Fluor Goat anti-rabbit 594 (Invitrogen, 1:2,000) were added for 1 h at room temperature. Finally, several washes with PBS were performed and the slides were mounted with Mowiol, with or without DAPI to visualize the cell nuclei (according to the assay). A Carl Zeiss Axio Imager. M2 Microscope (ZEISS Microscopy, Germany) with a Plan-Apochromat 63x/1.40 Oil M27 objective and a 2.8-megapixel monochrome Zeiss 503 digital video camera was used.

### Determination of TgRAD51 foci in response to different treatments

For intracellular parasites, hTERT monolayers were cultured in 24-well plates and infected with 3×10⁵ RHΔKU80TIR1 or RAD51HA-AID tachyzoites per well. After 16 hours of infection, cultures were treated with 1 mM hydroxyurea (HU, Sigma Aldrich), 4 µM Camptothecin (CPT, Sigma-Aldrich) or DMSO for 4 hours. Stock solution of 10 mM CPT was made using dimethylsulfoxide (DMSO). For extracellular parasites, tachyzoites were directly collected, centrifuged, and resuspended in 100 µl of 4% PAF per well for 20 minutes. Immunofluorescence staining was followed as described above by using a rabbit anti-SAG1 antibody (1:2,000), rabbit anti-TgHsp90 antibody (1:500) and a mouse anti-HA antibody (1:200).

### Western blot

Freshly purified RAD51^HA-AID^ tachyzoites were centrifuged and resuspended in NuPAGE lysis buffer and sonicated prior to resolution on SDS-PAGE and transferred to a nitrocellulose membrane. HA epitopes were detected using anti-HA mouse antibody at a 1:200 dilution followed by secondary probing with horseradish peroxidase (HRP)-conjugated goat anti-rat antibody (GE Healthcare) at a 1:2,000 dilution. To ensure equal loading of samples, we also probed with rabbit anti-SAG1 at a 1:5,000 dilution followed by HRP-conjugated donkey anti-rabbit antibody (GE Healthcare) at a 1:2,000 dilution.

### Cell viability analysis

MTT (Sigma, M2128-1G) (3-(4,5-dimethylthiazol-2-yl)-2,5-diphenyl-tetrazolium bromide) assay was used to assess cell viability in response to 3-benzyl-2-[(E)-2-pyridin-3-ylethenyl]quinazolin-4-one (B02, Sigma Aldrich) treatment. In this study, hTERT cells were cultured in 96-well plates until they reached 80% confluence, then treated with different concentrations of B02 (0 to 50 µM). Stock solution of 25 mM B02 was prepared using DMSO. After 72 hours of incubation cells were washed twice with PBS and the medium was replaced with 100 µl of MTT solution (0.5 mg/ml in PBS). The MTT compound is reduced by living cells, producing a violet-colored product (formazan), while dead cells cannot carry out this conversion. Then, the formazan crystals were solubilized with DMSO and the absorbance at 540 nm was measured on Synergy H1 Hybrid Multi-Mode microplate reader (Synergy H1), indicating the number of viable cells present.

### IC_50_ value of B02

hTERT cells (2×10^4^ per well) were cultured in 96-well plates for 24 h prior to the addition of 10,000 freshly lysed-out RH-RFP tachyzoites per well. Half of the plate was inoculated with tachyzoites, meanwhile, the rest of the plate was used as control (host cells + different concentrations of the drugs tested). After infection, the plate was incubated on ice for 10 min and then placed at 37 °C for 2 h to allow invasion to proceed. The medium was then replaced with fresh medium containing different concentrations of B02 or the vehicle (DMSO) and incubated for 72 h at 37°C. The following concentrations were tested for B02: 15 µM, 7.5 µM, 3.75 µM, 1.875 µM, 0.9375 µM and 0.45875 µM. Fluorescence values, expressed as relative fluorescence intensity units or RFU (excitation 544 nm; emission 590 nm) were obtained by reading from the bottom of the plate using the BioTek Synergy H1 microplate reader with an auto gain setup. For the half-maximal inhibitory concentration (IC_50_) calculations, the fluorescence value for each concentration was obtained by subtracting the fluorescence of the non-infected cells from that of infected cells. The data were then normalized to the fluorescence value obtained for the vehicle (100% fluorescence). Finally, the mean fluorescence percentage (% Mean RFU) was plotted against the logarithm of the tested concentration. IC_50_ values were obtained by nonlinear regression analysis of the data, using GraphPad Prism 8 software (GraphPad Software, Inc). Three independent experiments were performed.

### Replication assay

hTERT monolayers were infected with 10^5^ tachyzoites of the designated strain per well. RAD51^HA-AID^ parasites were treated with vehicle or 500 μM indole-3-acetic acid (IAA) overnight to knockdown TgRAD51. IAA stock solution was prepared by dissolving IAA in ethanol. After 1-hour incubation, cells were washed with PBS and incubated for 20 h with DMSO (control). Next day, cells were fixed with 100% methanol for 7 min at -20◦C and immunofluorescence staining (IFA) was performed. The number of tachyzoites per parasitophorous vacuole (PV) in 100 randomly selected fields was counted. Data is presented as the mean number of tachyzoites per PV. Statistical analysis was performed by GraphPad Prism 8, ANOVA, multiple comparisons. The effect of B02 on *T. gondii* replication was analyzed in a similar way. In this case, *T. gondii* lines RHΔKU80 or RHΔHXGPRT were incubated with 5 µM B02 overnight.

### Cell cycle analysis

hTERT cells were grown in 6-well plates and when confluent they were infected with 2x10^6^ RHΔHXGPRT tachyzoites per well. They were treated with different concentrations of B02 (10, 5 or 2.5 μM), hydroxyurea (HU 4 mM), or 0.1% v/v DMSO for 6 hours. Subsequently, the plates were washed with PBS and cells scrapped, collected and intracellular tachyzoites obtained after forcing through different sized needles. Tachyzoites were filtered using a 3 µm polycarbonate membrane (Gamafil). Purified tachyzoites were centrifuged at 2,000 RPM for 10 minutes, washed with PBS, and fixed with 70% ethanol for 24 hours at -20 °C. Samples were then centrifuged and washed with PBS supplemented with 2% FBS. Then tachyzoites were resuspended in 1 ml PBS supplemented with 180 µg/ml RNase and incubated for 10 min at 37°C. Finally, they were incubated with propidium iodide (0.5 mg/ml) for 10 minutes before measuring on the BD FACS Calibur flow cytometer. Results were analyzed with FlowJo10 (BD Biosciences).

### Differentiation assay

hTERT cells were grown on coverslips in 24-well plates and when confluent they were infected with 1x10^4^ ME49 tachyzoites, then placed at 37°C for 2 h to allow invasion. After washing with PBS, the

intracellular tachyzoites were incubated with fresh medium (pH 7.0) containing different concentrations of B02 (5µM or 10µM) or vehicle for 96 h at 37°C with CO_2_. For negative control (C-), intracellular ME49 parasites were incubated with DMEM at pH 7.0 with CO_2_. For positive control (C+), intracellular ME49 parasites were incubated at pH 8.1 without CO_2_ 5% to induce bradyzoite differentiation. *Dolichos biflorus*-agglutinin (1:200) from Vector Laboratories (FL-1031) was applied for 1 hour at room temperature to detect cyst walls. Anti-TgHSP90 rabbit antibody (1:500) and anti-SAG1 rabbit (1:2,000) were incubated 1 hour at room temperature. The immunofluorescence was performed as explained before. The number of cysts was counted manually in each field using the ImageJ application software 1.53t (Fiji) to confirm the positive DLB mark (20 fields per treatment were analyzed). The diameter of the cyst was measured using the straight tool by drawing a line corresponding to the diameter of each DBL+ cyst. Each photo was calibrated to the corresponding scale.

### Drugs combination

hTERT cells were cultured in 96-well plates for 24 h prior to the addition of 10,000 freshly lysed-out RH-RFP tachyzoites per well. Half of the plate was inoculated with tachyzoites, meanwhile, the rest of the plate was used as control (host cells + different concentrations of the drugs tested). After infection, the plate was incubated on ice for 10 min and then placed at 37 °C for 2 h to allow invasion. The medium was changed to add different concentrations of the drugs or vehicle (DMSO) and incubated to complete 72 h at 37°C. The following concentrations were tested: 2.5 µM B02; 2.5 µM CPT, 0.1 mM HU; 2.5 µM B02 + 2.5 µM CPT and 2.5 µM B02 5 + 0.1 mM HU. Fluorescence values (excitation 544 nm; emission 590 nm) were obtained by reading from the bottom of the plate using a Synergy H1 Hybrid Multi-Mode plate reader (BioTek) with an auto gain setup. Data analysis was performed as described in section 2.6. Three independent experiments were performed.

### Statistics

Data variations were analyzed with GraphPad Prism 8 software, using one-way ANOVA and Tukey’s multiple comparisons test (*p ≤ 0.05, **p ≤ 0.01, ***p ≤ 0.001, ****p < 0.0001) comparison test according to the assay.

## CRediT authorship contribution statement

**Ana M. Saldarriaga Cartagena**: Methodology, Investigation, Formal analysis, Writing – review & editing. **Ayelén Aparicio Arias:** Methodology, Investigation, Formal analysis. **Constanza Cristaldi**: Methodology, Investigation. **Agustina Ganuza**: Methodology. **M. Micaela Gonzalez**: Supervision, Methodology, Writing – review & editing. **Maria Corvi:** Supervision, Methodology, Writing – review & editing. **William J. Sullivan, Jr:** Writing – original draft, Validation, Funding acquisition, Formal analysis, Conceptualization. **Laura Vanagas:** Writing – original draft, Validation, Funding acquisition, Formal analysis, Conceptualization. **Sergio O. Angel:** Writing – original draft, Validation, Funding acquisition, Formal analysis, Conceptualization

## Ethical approval

Does not correspond

## Consent for publication

Does not correspond

## Availability of data and materials

The authors confirm that the data supporting the findings of this study are available within the article and its supplementary materials. The data used and analyzed during the current study are also available from the corresponding author upon reasonable request.

## Funding

This work was supported by the Ministerio Nacional de Ciencia y Tecnología (MINCyT): PICT 2015 1288 (S.O.A. and M.M.C.), PICT 2021 0169 (L.V.), Consejo Nacional de Investigaciones Científicas y Tecnológicas (CONICET): PIP 11220150100145CO and 11220210100572CO (S.O.A., L.V., and M.M.C.) and by National Institute of Health: NIH-NIAID R01AI129807 (S.O.A. and W.J.S.).

## Declaration of competing interest

The authors declare that they have no known competing financial interests or personal relationships that could have appeared to influence the work reported in this paper.

## Author statement

All authors read and approved the submitted manuscript.

## Acknowledgments

M.M. Gonzalez, M.M. Corvi, L.Vanagas and S.O. Angel are members of Consejo Nacional de Investigaciones Científicas y Técnicas (CONICET) and Professors at Universidad Nacional General San Martin (UNSAM). A.M. Saldarriaga Cartagena and C. Cristaldi are doctoral fellow of CONICET. A. Aparicio is a Fellow of Fondo para la Investigación Científica y Tecnológica (FONCyT). A. Ganuza is a technical staff of CONICET.

## Notes

### Competing Interest Statement

The authors have declared no competing interest.

