## Supplemental figures for "*Toxoplasma gondii* RAD51 recombinase is required to overcome DNA replication stress and its inactivation leads to bradyzoite differentiation"

SUPPLEMENTAL FIGURE LEGENDS

Fig. S1. Generation of RAD51HA-AID line. Positive clones of RHΔKU80TIR1 *T. gondii* line that integrated the RAD51-AID-HA tag were validated by PCR of both 5' -end and 3' -end. To amplify the RAD51 gene we use this pair of primers: forward: CGCCATCGGCGAAGGAGGCATCGGCGACTACGAAGACAAccgctagcaagggctcggg (P1), and reverse: ctgcatccgtgtagctctgtgactttgagcctgttggaacaaaagctggagctccac (P4). The primers sequence for validation includes the donor primers and an extra pair: forward: gaacatcctcaacaaggaac (P3) and reverse: gccaccgcttgatttttggca (P2). Negative parasites and parental RHΔKU80TIR1 amplify a 396bp band using P1P4 primers, positive clones amplify three bands to check the correct insertion of the AID-HA tag: P1P2=536bp, P3P4=1322bp, P1P4=5572bp.

Fig. S2. Identification of RAD51-like genes from *T. gondii*. *T. gondii* genes associated with recombinases were detected in ToxoDB by searching by name and sequence. Cell cycle expression (transcriptome section) analyses and fitness phenotype data were obtained from ToxoDB. In the case of TGME49_313710, its expression was analyzed by the RNA-seq data of Waldman et al (transcriptome section). Domains and motifs were analyzed by Motifscan web page (https://myhits.sib.swiss/cgi-bin/motif_scan).

Fig. S3. The parental line does not show RAD51 foci. Intra- and extracellular tachyzoites from the RHΔKU80TIR1 parental line were incubated with the anti-HA antibody that labels RAD51-HA-AID foci. No positive foci labeling was observed in any of the cases. The anti-SAG1 antibody was used to label the parasite. Nuclei are stained blue with DAPI.

Fig. S4. RAD51 foci increase with HU. Intracellular tachyzoites from the generated RAD51$^{-HA-AID}$ line were incubated with 5µM CPT or 1mM HU overnight. DMSO 0.1% was used as a control. Rad51 foci are observed with the anti-HA antibody in green. Nuclei are stained blue with DAPI.

Fig. S5. Alignment of RAD51 amino acid sequences. The amino acid sequences of TgRAD51 (TGME49_272900), *P. falciparum* PfRAD51 (PlasmoDB gene ID PF3D7_1107400), *E. histolytica* EhRAD51 (GenBank ID XM_648984) and Human RAD51 (HuRAD51, uniprot Q06609) were aligned by clustal W.

FigS1

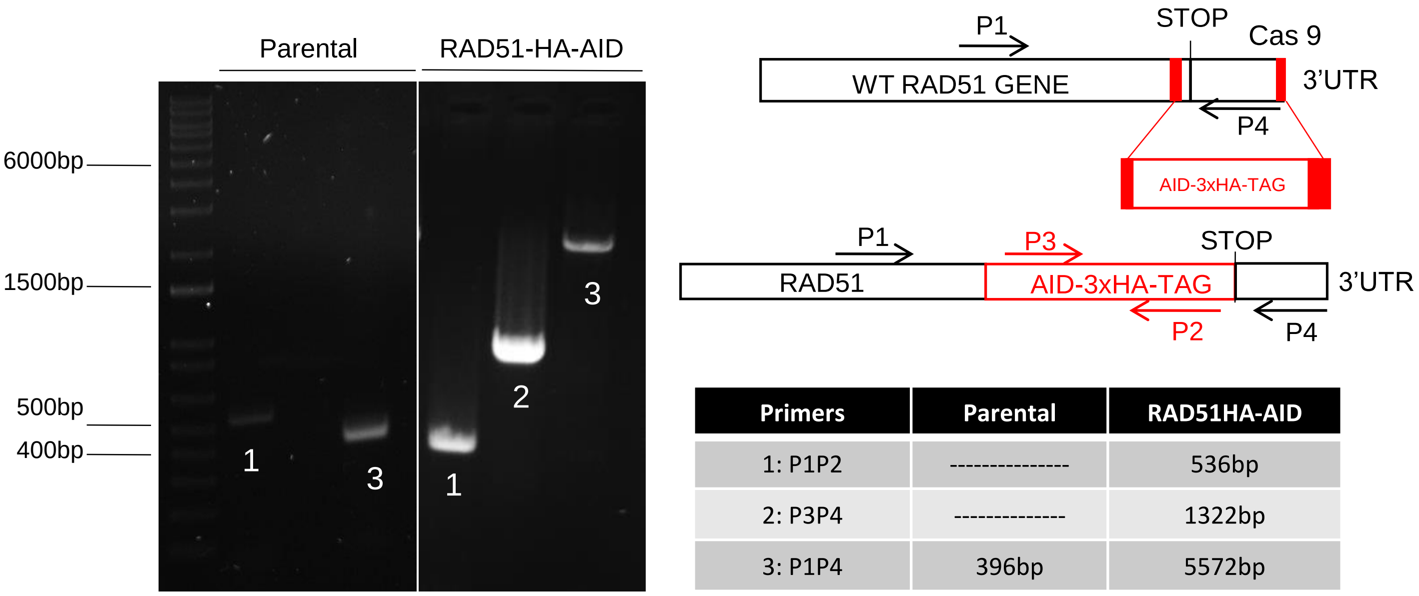


| Primers | Parental | RAD51HA-AID |
|---|---|---|
| 1: P1P2 | --------------- | 536bp |
| 2: P3P4 | -------------- | 1322bp |
| 3: P1P4 | 396bp | 5572bp |

Fig S2

TGME49_272900, DNA repair protein RAD51 — Phenotype: -0.34

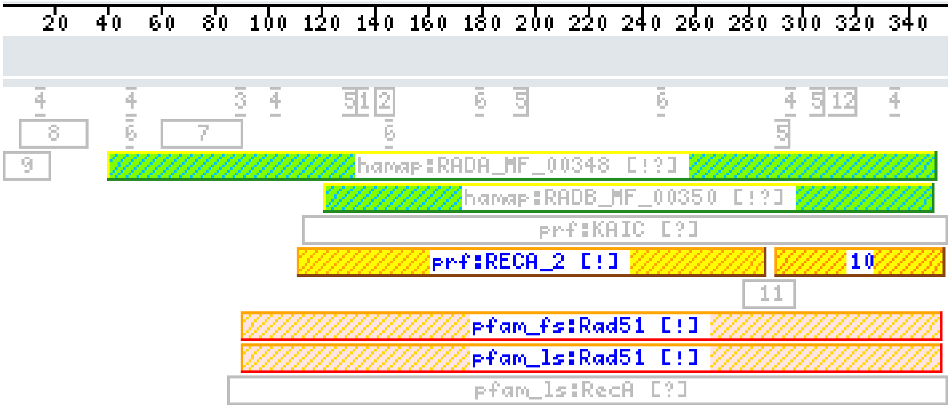


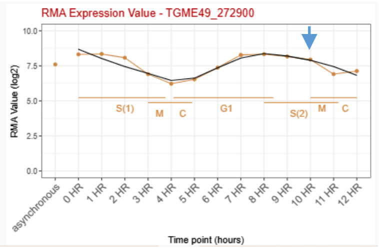


TGME49_321430, DNA repair protein recA — Phenotype: -0.35

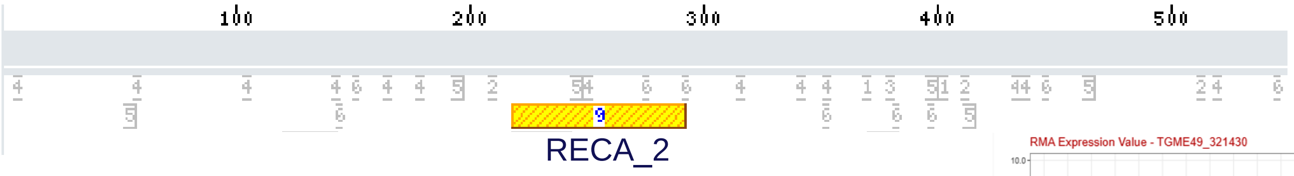


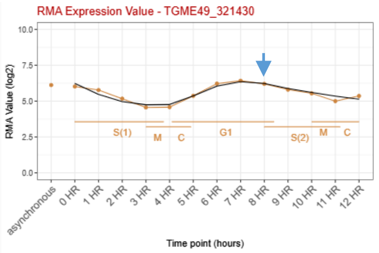


TGME49_313710, hypothetical protein — Phenotype: -0.77

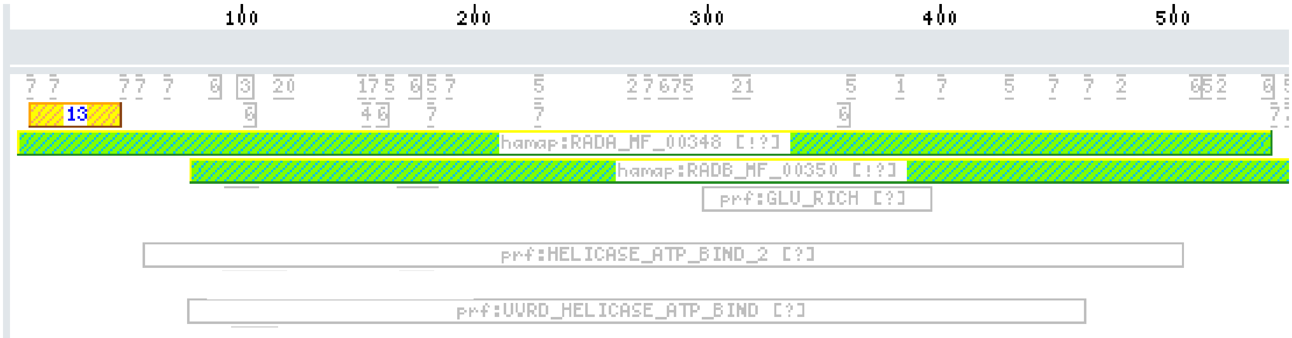


TGME49_216400 meiotic recombination protein DMC1 — Phenotype: -0.1

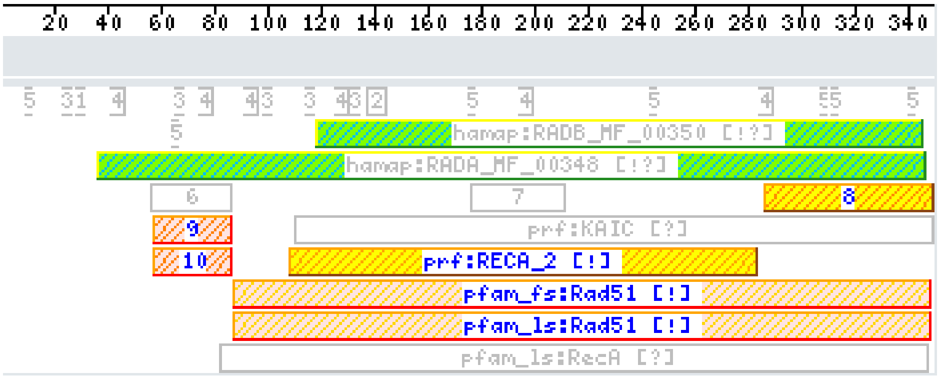


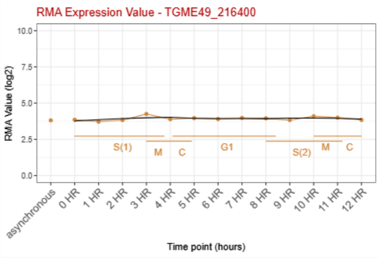

Fig. S3

#### Parental

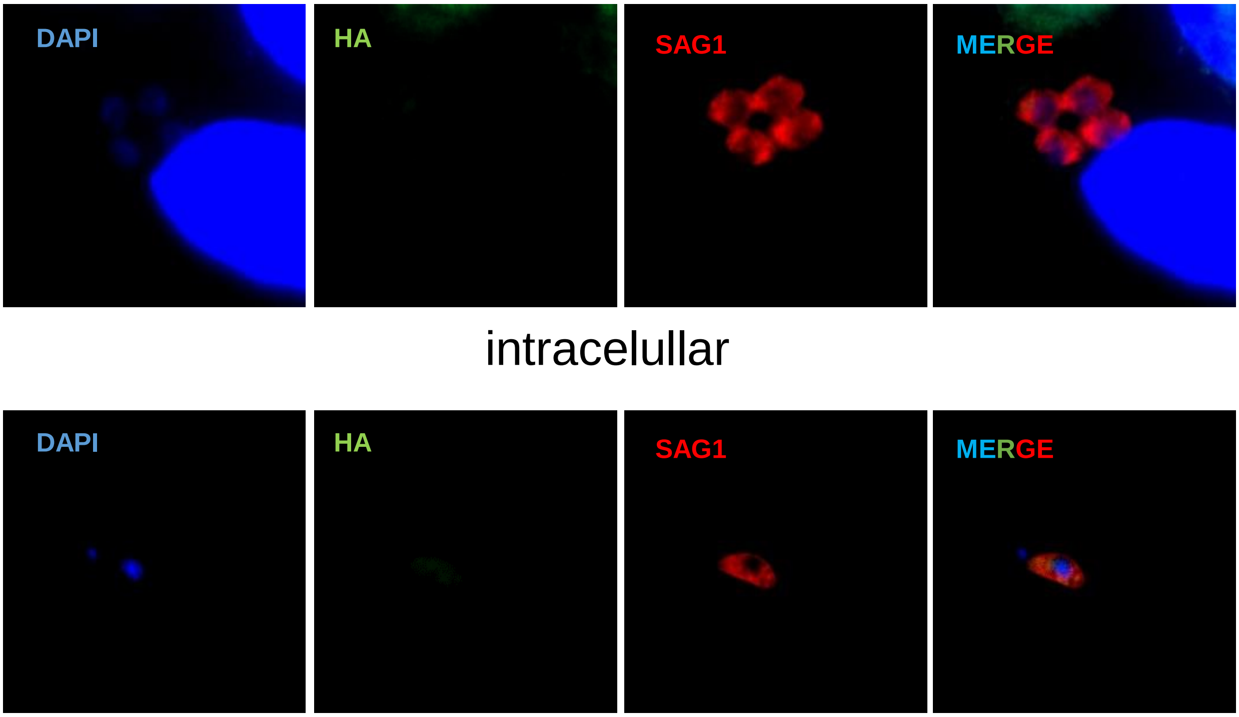


intracelullar

extracelullar

Fig. S4

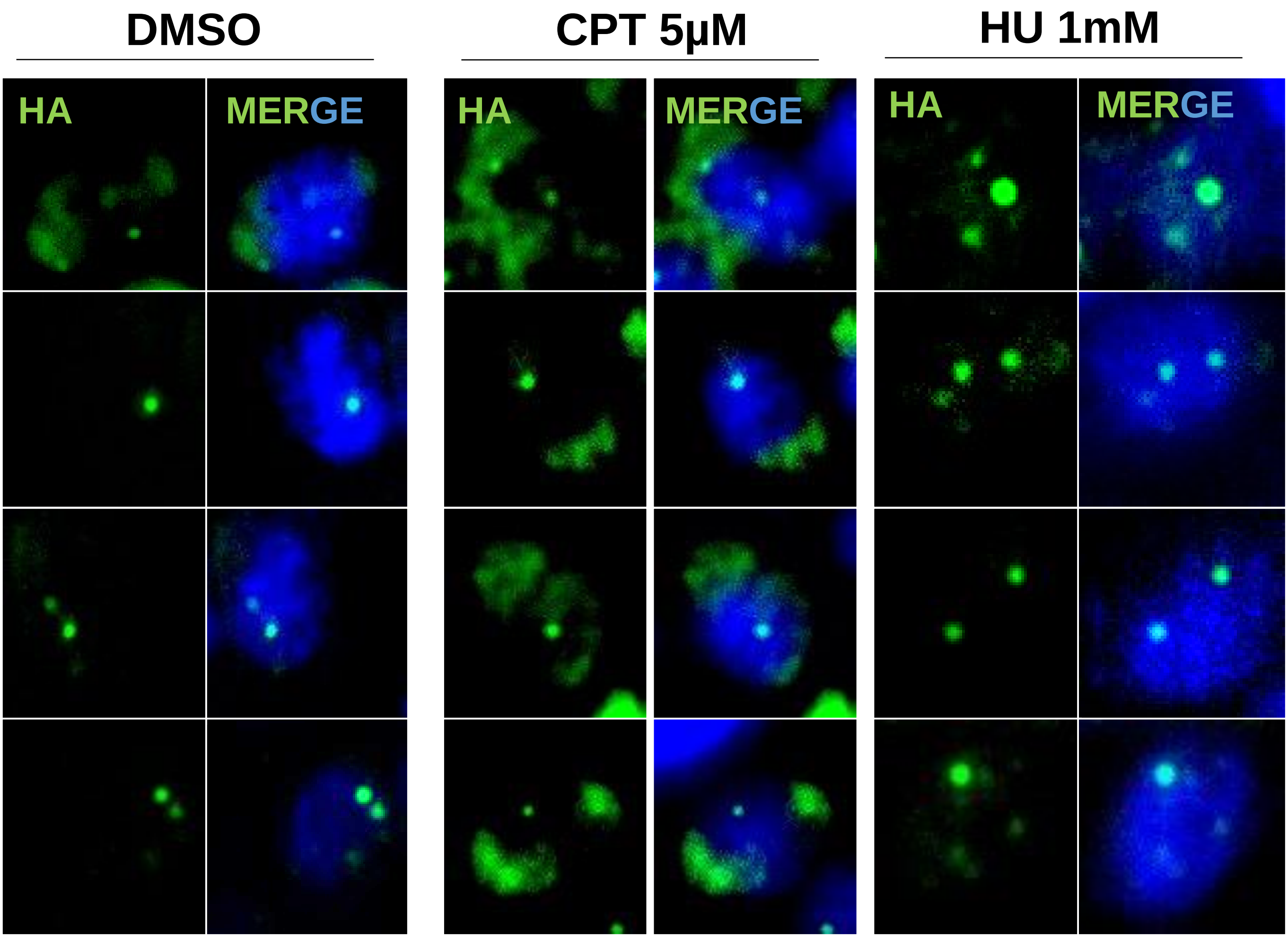

### Fig S5

```
                  10        20        30        40        50        60        70        80
TgRAD51  MSAVSLQQSRAASVQESEP-----------QQQQAQQLAEEVQSGPLKLEHLL-AKGFTKRDLELLKDAGYQTVECIAFA
PfRAD51  MKQANTKEDKSQKISNSST-----------IDEIEE---EQLYTGPLKIEQLL-AKGFVKRDLELLKEGGLQTVECVAYA
HuRAD51  -MAMQMQ-------LEAN----------------ADTSVEEESFGPQPISRLE-QCGINANDVKKLEEAGFHTVEAVAYA
EhRAD51  MSAKQIQRGSEEELLESEENLKEKEKVNEEEHEEEEEEEEEEENAAVPIRKLIGLAGITEGDCKKLEEAGFFTVQSIAFT

                  90       100       110       120       130       140       150       160
TgRAD51  PVKNLVAVKGLSEQKVEKLKKASKELCNLGFCSAQEYLEARENLIRFTTGSVQLDSLLKGGIETGNLTELFGEFRTGKTQ
PfRAD51  PMRTLCAIKGISEQKAEKLKKACKELCNSGFCNAIDYHDARQNLIKFTTGSKQLDALLKGGIETGGITELFGEFRTGKSQ
HuRAD51  PKKELINIKGISEAKADKILAEAAKLVPMGFTTATEFHQRRSEIIQITTGSKELDKLLQGGIETGSITEMFGEFRTGKTQ
EhRAD51  PKKQLITIKGISDAKADKLLAESSKIVPLAFTNAAELNNLRKETIRITTGSRELDKLLCGGFETGSITELFGEFRTGKTQ

                 170       180       190       200       210       220       230       240
TgRAD51  LCHTLAVTCQLPIEQAGGEGKCLWIDTEGTFRPERIVSIAKRFGLNANDCLDNVAYARAYNCDHQMELLMEASAMMAESR
PfRAD51  LCHTLAITCQLPIEQSGGEGKCLWIDTEGTFRPERIVAIAKRYGLHPTDCLNNIAYAKAYNCDHQTELLIDASAMMADAR
HuRAD51  ICHTLAVTCQLPIDRGGGEGKAMYIDTEGTFRPERLLAVAERYGLSGSDVLDNVAYARAFNTDHQTQLLYQASAMMVESR
EhRAD51  LCHQLCVTCQLGIENGGTEGRAIYIDTEGTFRPERLTQIAEKYGLNSEEALNNVAVARAHNTEHQMQLLQMASGLMAKER

                 250       260       270       280       290       300       310       320
TgRAD51  FALLIVDSATALYRSEYTGRGELASRQTHLCRFLRCLQRIADTYGVAVVVSNQVVAKVDNMGGMFSGNEKLPIGGNIMAH
PfRAD51  FALLIVDSATALYRSEYIGRGELANRQSHLCRFLRGLQRIADIYGVAVIITNQVVAKVDAMS-MFGGHEKIPIGGNIIAH
HuRAD51  YALLIVDSATALYRTDYSGRGELSARQMHLARFLRMLLRLADEFGVAVVITNQVVAQVDG-AAMFAADPKKPIGGNIIAH
EhRAD51  YGLLIIDSATALYRTDYSGRGELASRQMHLAKFLRALQRIADEFSVAVVLTNQVVAQVDGQASMFGGDTKKPVGGNIIAH

                 330       340       350       360
TgRAD51  ASQTRLYLRKGRGESRICKIYDSPSLAEGEAVFAIGEGGIGDYEDN
PfRAD51  ASQTRLYLRKGRGESRICKIYDSPVLPEGEAVFAITEGGIADYEEK
HuRAD51  ASTTRLYLRKGRGETRICKIYDSPCLPEAEAMFAINADGVGDAKD-
EhRAD51  ASTTRLYLRKGKGEARICKVYDSPCLPESEASFAITTNGIEDVKDD
```
